# Loss of KDM6A-mediated genomic instability and metabolic reprogramming differentially regulates responses to immune checkpoint therapy and chemotherapy in bladder cancer

**DOI:** 10.1101/2024.10.31.621396

**Authors:** Pratishtha Singh, Deblina Raychaudhuri, Bidisha Chakraborty, Swadhin Meher, Aminah J. Tannir, Anurag Majumdar, Jessalyn Hawkins, Yun Xiong, Philip Lorenzi, Padmanee Sharma, Patrick Pilié, Sangeeta Goswami

**Affiliations:** Department of Immunology, The University of Texas MD Anderson Cancer Center, Houston, Texas; Department of Bioinformatics and Computational Biology, Division, The University of Texas MD Anderson Cancer Center, Houston, Texas; Department of Genitourinary Medical Oncology, The University of Texas MD Anderson Cancer Center, Houston, Texas; Immunotherapy Platform, The University of Texas MD Anderson Cancer Center, Houston, Texas; James P. Allison Institute, The University of Texas MD Anderson Cancer Center, Houston, Texas

## Abstract

Mutations in genes encoding critical epigenetic regulators are frequently noted in bladder cancer, however, the impact of these mutations on therapeutic efficacy is unclear. One of the most common driver mutations in bladder cancer occurs in the *KDM6A* gene, which encodes a histone demethylase that promotes gene transcription. Retrospective analyses of patients with bladder cancer demonstrated that *KDM6A* mutations correlate with improved overall survival (OS) with immune checkpoint therapy (ICT), while they are associated with lower OS in patients undergoing cisplatin-based chemotherapy. Mechanistic studies utilizing CRISPR-Cas9 mediated deletion of *Kdm6a* showed reduced expression of DNA mismatch repair (MMR) and DNA double-stranded base repair (DSBR) genes in tumor cells with improved response to anti-PD-1 therapy and attenuated sensitivity to cisplatin-based chemotherapy in preclinical models of bladder cancer. Additionally, the loss of *Kdm6a*-mediated reduction in glycolysis and intratumoral lactate accumulation impaired histone 3 lysine 9 lactylation (H3K9la) and histone 3 lysine 18 lactylation (H3K18la) in Tregs with concurrent decrease in the expression of key genes including *Foxp3, Tgfb* and *Pdcd1* and their immune-suppressive function. Further, reduced expansion of PD-1^hi^ Tregs improved the ratio of cytotoxic T cells to Tregs and response to anti-PD-1 therapy in *Kdm6a* deficient tumor-bearing mice. Collectively, this study provided key insights into the role of KDM6A-mediated epigenetic regulation of DNA repair and metabolic reprogramming which potentially govern response to chemotherapy and ICT thus highlighting the utility of *KDM6A* mutation status for patient stratification and development of personalized treatment algorithms.

## Introduction

Bladder cancer is the sixth most commonly occurring cancer in the United States, with the five-year overall survival (OS) for advanced bladder cancer being less than 10% (1). Cisplatin-based chemotherapy has been a mainstay of therapy for many decades; however, the advent of immune checkpoint therapy (ICT) and targeted therapy has changed the current landscape of treatment for bladder cancer (2, 3). While the availability of different therapeutic agents such as chemotherapy, ICT and targeted therapy, either as single agents or in combination, has significantly improved outcomes, there remains a lack of biological insight into selecting and sequencing these therapies based on patient attributes to develop a personalized treatment algorithm.

Genes encoding key epigenetic regulators are frequently mutated in bladder cancer (4, 5). These epigenetic factors orchestrate gene expression, impacting multiple pathways governing cellular phenotype and function (6, 7). While the impact of mutations in epigenetic factors on initiation and progression of bladder tumorigenesis has been studied (8–10), the role of these mutations in regulating response to therapeutic agents, including chemotherapy and ICT, remains largely unexplored. Lysine Demethylase 6A (*KDM6A*) is a commonly mutated gene in bladder cancer, with approximately 26% of patients with muscle-invasive bladder cancer harbor *KDM6A* mutations (4, 11). KDM6A catalyzes the demethylation of H3K27 di- and tri-methyl (H3K27me2/me3) marks which are repressive histone post-translational modifications (PTMs), thereby promoting transcriptional activation (12, 13). Here, we investigated how inactivating mutations in *KDM6A* regulate responses to therapeutic perturbations including ICT and chemotherapy in bladder cancer.

Our retrospective analyses of advanced bladder cancer cohorts(14, 15) demonstrated improved OS of patients harboring *KDM6A* mutations in response to ICT compared to patients who do not have *KDM6A* mutations. Conversely, analysis of patient cohorts with resectable and advanced bladder cancer receiving cisplatin-based chemotherapy (16, 17) showed that patients with *KDM6A* mutations have decreased OS with cisplatin-based chemotherapy, indicating distinct roles of KDM6A in modulating therapeutic responses in bladder cancer. We also noted increased tumor mutation burden (TMB) in patients harboring *KDM6A* mutation. Importantly, a preclinical bladder tumor model harboring *Kdm6a* deletion (sgKdm6a) in a bladder cancer cell line demonstrated that while the sgKdm6a tumor-bearing mice have improved response to anti-PD-1 therapy, they exhibited decreased sensitivity to cisplatin-based chemotherapy, corroborating the findings from the clinical trials.

Mechanistic studies elucidated that KDM6A directly binds to genes associated with MMR and DSBR pathways and the loss of KDM6A resulted in the downregulation of MMR and DSBR-associated genes. Attenuated MMR activity and accumulation of DNA damage following the loss of KDM6A could partly account for the improved response to ICT and the decreased sensitivity to cisplatin. Additionally, we noted that the loss of KDM6A decreased glycolysis in the tumor cells with concomitant reduction in the lactate levels in the tumor microenvironment (TME). Importantly, we also observed reduced abundance of Tregs in sgKdm6a tumor-bearing mice with attenuated immune-suppressive function of the Tregs, possibly due to the lack of lactate in the TME as intratumoral Tregs utilize lactate as a fuel source to support their immune-suppressive functions (Watson et al., 2021). Further, we delineated a previously unknown pathway of lactate-mediated epigenetic regulation of Treg gene expression and immune-suppressive function. We demonstrated that the reduction in extracellular lactate impaired histone 3 lysine 9 lactylation (H3K9la) and histone 3 lysine 18 lactylation (H3K18la) in Tregs, attenuating transcription of genes including *Foxp3, Tgfb*, and *Pdcd1,* thus diminishing the immune-suppressive activity of Tregs. Additionally, reduced PD-1 expression on Tregs in sgKdm6a tumors decreased anti-PD-1 therapy mediated expansion of PD-1^hi^ Tregs thus improving the ratio of cytotoxic T cells to Tregs and response to anti-PD-1 therapy.

Overall, this study provided critical mechanistic insights into the role of KDM6A in modulating tumor cell-intrinsic pathways such as DNA damage response (DDR) pathways, which subsequently govern the response to different therapeutic perturbations including ICT and chemotherapy. Further, this study delineated how the loss of KDM6A suppresses glycolysis and lactate production in tumor cells, which subsequently attenuates histone lactylation in intratumoral Tregs thereby epigenetically regulating immune-suppressive functions of Tregs. Together, these findings underscored the role of *KDM6A* mutations in regulating therapeutic sensitivity in bladder cancer thus guiding patient selection for personalized treatment strategies.

## Results

### Inactivating *KDM6A* mutations improve OS with ICT while it reduces OS with cisplatin in bladder cancer

To garner insight into how inactivating mutations in *KDM6A* impact responses to different therapeutic perturbations in bladder cancer, we performed retrospective analysis of data from patients receiving ICT and cisplatin-based chemotherapy. Analyses of the IMVigor210 (N = 275) cohort where patients with metastatic bladder cancer received anti-PD-L1 therapy (14) demonstrated that patients harboring the *KDM6A* mutation (KDM6A-Mut) had significantly improved overall survival (OS) in response to anti-PD-L1 therapy compared to patients without *KDM6A* mutation (KDM6A-WT) (Fig. 1A). Additionally, analyses of another retrospective cohort (MSK_2018) (15) of patients with advanced bladder cancer receiving ICT also showed improved OS in KDM6A-Mut patients (Fig. S1A). Importantly, analysis of patient cohorts (IMVigor210, N=275 and TCGA, N=136) (4, 14), revealed that patients with the *KDM6A* mutation had higher TMB (Fig. 1B). Since higher TMB is frequently driven by mutations in genes encoding enzymes involved in DNA MMR (18), we investigated the frequency of MMR gene mutations in the *KDM6A* mutated patients. However, we did not note any cooccurrence of the *KDM6A* mutation with mutations in the genes associated with the MMR pathway in these patients (Fig. 1C). Notably, analyses of the HCRN dataset (19) demonstrated decreased expression of several critical genes involved in the MMR pathway including *MSH2* and *MSH6* in patients harboring the *KDM6A* mutation indicating an attenuated MMR machinery in these patients (Fig. S1B). Thus, highlighting a distinct KDM6A-mediated pathway regulating genes associated with the MMR machinery and TMB. To confirm the association of inactivating KDM6A mutations with improved response to anti-PD-1 therapy, we generated CRISPR-Cas9-mediated *Kdm6a*-deleted murine MB49 bladder cancer cell line (sgKdm6a) (Fig. S1C). Consistent with the findings from the clinical cohorts, deletion of *Kdm6a* in murine bladder cancer cell line attenuated tumor growth following anti-PD-1 therapy in tumor-bearing mice (Fig. 1D).

**Figure 1.**
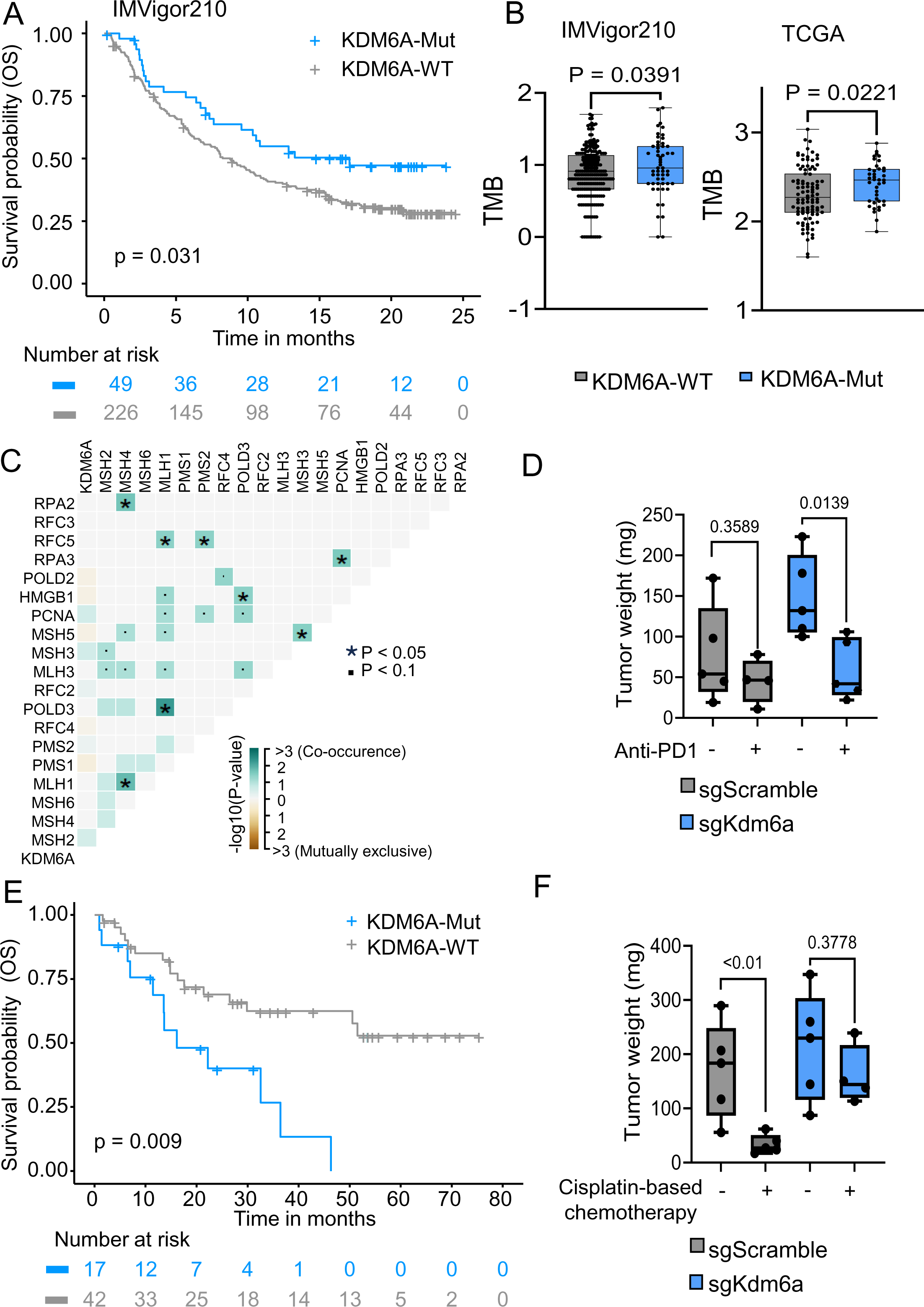
Inactivating KDM6A mutations improve OS with ICT while it reduces OS with cisplatin in bladder cancer. **(A)** Kaplan-Meier plot displaying the overall survival (OS) of advanced bladder cancer patients with *KDM6A* mutation (KDM6A-Mut) and without KDM6A mutation (KDM6A-WT) treated with anti-PD-1/L-1 therapy in IMVigor210 clinical trials. (n=275 patients, KDM6A-Mut=49, KDM6A-WT=226) Two-tailed Log-rank test was performed. **(B)** Box-and-whisker plots representing tumor mutation burden (TMB) in KDM6A-WT and KDM6A-Mut patients, from two patient cohorts. (IMVigor210, n=275 patients; TCGA BLCA, n=136 patients) Two-sided Wilcoxon test was performed. **(C)** Somatic interaction plot depicting co-occurrence or mutual exclusivity of *KDM6A* mutations with mutations in mismatch repair pathway (MMR) genes in bladder cancer patients in the TCGA BLCA dataset harboring *KDM6A* mutation (n=412 patients). Two-tailed Pair wise Fisher’s exact test was performed. **(D)** Box-and-whisker plot showing the tumor weights obtained from sgScramble and sgKdm6a tumor bearing female mice treated with and without anti-PD-1. (n= 4 female mice per group) Two-tailed Student’s t-test was performed. **(E)** Kaplan-Meier plots demonstrating OS of KDM6A-WT and KDM6A-Mut patients who recieved platinum-based chemotherapy (n=59 patients, KDM6A-Mut =17, KDM6A-WT=42). Two-tailed Log-rank test was performed. **(F)** Box-and-whisker plot illustrating the tumor weights derived from sgScramble and sgKdm6a tumor harboring female mice treated with and without 2.5 mg/kg Gemcitabine plus 6 mg/kg cisplatin (n= 4 female mice per group). Two-tailed Student’s t-test was performed. For all the box-and-whisker plots in Figure 1, the centre line marks the median, the edges of the box represent the interquartile (25th–75th) percentile and the whiskers represent minimum-maximum values.

Next, to delineate the role of *KDM6A* mutation in regulating the response to cisplatin-based chemotherapy, we analyzed OS of patients with resectable and advanced bladder cancer undergoing cisplatin-based chemotherapy (16, 17). Contrary to the improved OS noted in response to ICT in patients harboring the *KDM6A* mutation, we observed a reduction in OS with platinum-based chemotherapy in patients with the *KDM6A* mutation compared to patients without the *KDM6A* mutation (Fig. 1E). Consistently, mice harboring sgKdm6a tumors showed resistance to cisplatin-based chemotherapy compared to mice harboring control (sgScramble) tumors (Fig. 1F). To further investigate the differential impact of cisplatin on sgScramble versus sgKdm6a cell lines, we treated the cell lines with cisplatin *in-vitro*. We observed significantly higher cisplatin-induced cytotoxicity in sgScramble cells compared to sgKdm6a cells (Fig. S1D). Additionally, sgKdm6a cells demonstrated higher migration and spheroid forming potential in response to cisplatin, compared to sgScramble cells (Fig. S1E, S1F), indicating reduced sensitivity of *Kdm6a*-deleted murine bladder cancer cell line to cisplatin.

Together, retrospective analyses of patient cohorts and preclinical models demonstrated that KDM6A plays a differential role in modulating the response to ICT and cisplatin-based chemotherapy. While loss of *Kdm6a* improves response to ICT, it lowers sensitivity to cisplatin. Therefore, *KDM6A* mutation status could potentially be used to aid patient stratification for cisplatin-based chemotherapy and ICT.

### KDM6A directly regulates the DNA mismatch repair pathway in bladder cancer cells

Next, to investigate the mechanisms by which KDM6A regulates the expression of genes involved in the MMR pathway in bladder cancer, we performed chromatin immunoprecipitation sequencing (ChIP-seq) of sgScramble and sgKdm6a cells. We noted direct KDM6A binding to multiple MMR genes including *Msh2, Msh6,* and *Mlh1* in sgScramble cells (Fig. 2 A,B,D) with lack of binding of KDM6A to these genes following the loss of *Kdm6a* in sgKdm6a cells (Fig. 2 A,B,D). Importantly, we also observed corresponding attenuation of H3K4me3 enrichment in these genes in the absence of KDM6A (Fig. 2 A,B,Fig. S2A).

**Figure 2.**
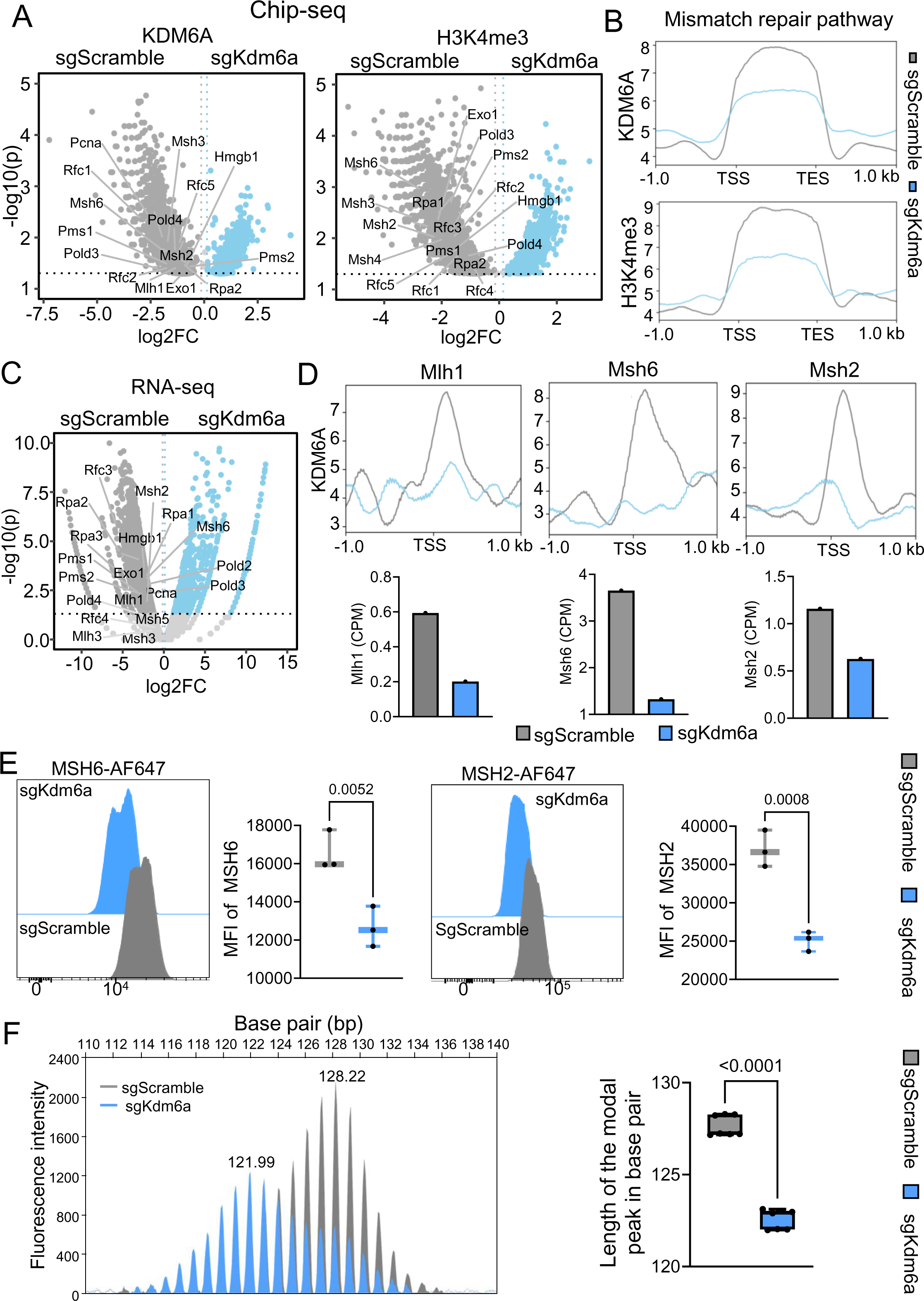
KDM6A directly regulates the DNA mismatch repair pathway in bladder cancer cells. **(A)** Volcano plots demonstrating the differential enrichment of KDM6A (Left Panel) and H3K4me3 (Right Panel) marks in the indicated genes involved in DNA MMR pathway between sgScramble and sgKdm6a cell lines, from ChIP-seq study. Volcano plots show the log2 ratio of fold change (log2FC) plotted against the Absolute Confidence, -log10 adjusted p-value (log10(p The P-values represent two-tailed probability distribution based on Bayesian model calculated using MAnorm. **(B)** Profile plots depicting the probability score of KDM6A (Top Panel) and H3K4me3 (Bottom Panel) binding at ±1kb regions from TSS and Transcription End Site (TES) for genes involved in DNA MMR pathway in sgScramble and sgKdm6a cells. **(C)** Volcano plot representing differential expression of the indicated genes involved in DNA MMR pathway between sgScramble and sgKdm6a cell lines from RNA-seq study. Volcano plots show the log2 ratio of fold change (log2FC) plotted against the Absolute Confidence, -log10 adjusted p-value (log10(p)). P-values were calculated with two tailed exact tests under a negative binomial distribution using EdgeR and adjusted with Benjamini–Hochberg method. **(D)** Profile plots (Top Panel) displaying the probability scores of KDM6A binding at promoter regions (TSS ±1kb) of *Mlh1*, *Msh6* and *Msh2* gene loci in sgScramble and sgKdm6a cell lines and count per million (CPM) bar plots (Bottom Panel) illustrating RNA expression levels of the corresponding genes. **(E)** Representative histograms (Left Panel) and box-and-whisker plots (Right Panel) showing the difference in Median Fluorescence Intensity (MFI) of MSH6 and MSH2 between sgScramble and sgKdm6a cells. (n=3 biologically independent samples per group) Two-tailed Student’s t-test was performed. **(F)** Representative electropherogram (Left Panel) derived from Fragment Fluorescent Length Analysis (FFLA) of *mBAT-64* microsatellite from sgScramble and sgKdm6a tumor cells and corresponding box-and-whisker plot (Right Panel) representing the length of the modal peak in base pair in sgScramble and sgKdm6a tumor cells. (n=7 female mice per group) Two-tailed Student’s t-test was performed. For all the box-and-whisker plots in Figure 2, the centre line marks the median, the edges of the box represent the interquartile (25th–75th) percentile and the whiskers represent minimum-maximum values.

We used RNA-sequencing (RNA-seq) based gene expression studies to confirm the downregulation of the MMR genes in sgKdm6a cells. RNA-seq demonstrated a concurrent reduction in the expression of the MMR genes in sgKdm6a cells (Fig. 2 C,D), mirroring the findings from the patient cohorts, suggesting KDM6A-and H3K4me3-mediated regulation of these genes. To confirm the reduction in MMR enzyme expression at the protein level, we performed flow cytometry. This analysis revealed decreased expression of MSH2 and MSH6 in sgKdm6a cells compared to sgScramble cells (Fig. 2E).

Disrupted MMR pathway is associated with microsatellite instability and an MSI-h phenotype (20, 21). Therefore, to determine the functional impact of the decreased expression of MMR genes in sgKdm6a tumor cells, we compared microsatellite instability (MSI) in sgScramble versus sgKdm6a MB49 tumors. We used fluorescent fragment length analysis (FFLA) to compare the MSI profiles of sgScramble (N=7) and sgKdm6a (N=7) MB49 tumors. FFLA involves labeling DNA fragments with fluorescent dyes and analyzing their sizes and distribution via capillary electrophoresis. Importantly, analysis of electropherogram peaks revealed a significant left shift of 5 nucleotides in the modal (tallest) peak in sgKdm6a tumors compared to sgScramble tumors indicating contraction of the *mBAT-64* microsatellite length and genetic instability (Fig. 2F). Cumulatively, these findings indicate that KDM6A modulates H3K4me3 enrichment at the promoter regions of DNA MMR genes thus regulating their expression and the microsatellite instability status in tumor cells.

The MMR-deficient, MSI-h phenotype has been associated with improved responses to ICT (22, 23) and reduced sensitivity to platinum-based chemotherapy (24–27). Therefore, attenuated MMR activity following the loss of KDM6A could account for improved response to ICT while decreasing sensitivity to cisplatin.

### Loss of KDM6A impairs the double-stranded break repair pathway

In addition to reduced expression of genes involved in the MMR pathway following the loss of KDM6A, we observed lower expression of genes involved in DSBR, including *EXO1* and *LIG1,* in bladder cancer patients harboring *KDM6A* mutations (19) (Fig. S3A). Importantly, we did not observe any cooccurrence of the *KDM6A* mutation with mutations in the DSBR genes in these patients suggesting an independent role of KDM6A in regulating the expression of DSBR genes (Fig. S3B).

Next, we analyzed the murine ChIP-seq data to investigate the impact of KDM6A loss on genes involved in DSBR. Our results showed a reduction in KDM6A binding in multiple genes involved in DSBR pathways including *Exo1* and *Lig3* in sgKdm6a cells indicating KDM6A-mediated regulation of these genes (Fig. 3A, C, Fig. S3C). We also observed a concurrent decrease in H3K4me3 enrichment across these genes (Fig. 3A, Fig. S3C, D). RNA-seq analysis also demonstrated a reduction in the expression of these DSBR genes in sgKdm6a MB49 cells, confirming the findings from the ChIP-seq data (Fig. 3B, C).

**Figure 3.**
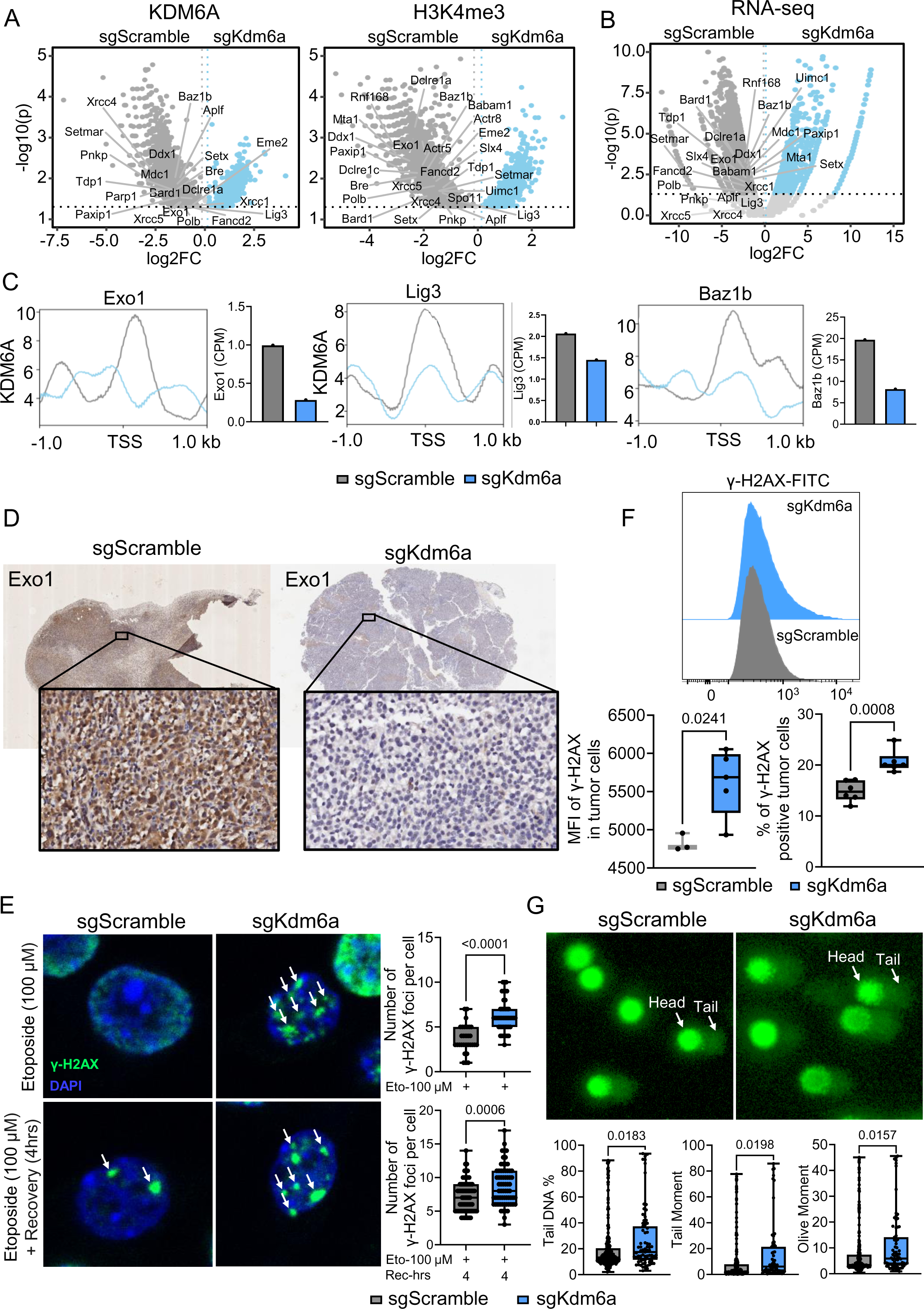
Loss of KDM6A impairs the double-stranded break repair pathway. **(A)** Volcano plots demonstrating the differential enrichment of KDM6A (Left Panel) and H3K4me3 (Right Panel) marks in the indicated genes involved in double-strand break repair (DSBR) pathway between sgScramble and sgKdm6a cell lines, from ChIP-seq data. Volcano plots show the log2 ratio of fold change (log2FC) plotted against the Absolute Confidence, -log10 adjusted p-value (log10(p)). The P-values represent two-tailed probability distribution based on Bayesian model calculated using MAnorm. **(B)** Volcano plot representing differential expression of the indicated genes involved in DSBR pathway between sgScramble and sgKdm6a cell lines from RNA-seq data. Volcano plots show the log2 ratio of fold change (log2FC) plotted against the Absolute Confidence, -log10 adjusted p-value (log10(p)). P-values were calculated with two tailed exact test under a negative binomial distribution using EdgeR, and adjusted with Benjamini–Hochberg method. **(C)** Profile plots depicting the probability scores of KDM6A binding at promoter regions (TSS ±1kb) of *Exo1*, *Lig3* and *Baz1b* gene loci in sgScramble and sgKdm6a cell lines and count per million (CPM) bar plots illustrating RNA expression levels of the corresponding genes. **(D)** Representative figures showing the immunohistochemical (IHC) staining for EXO1 expression (brown) in tissue sections derived from sgScramble and sgKdm6a tumors. Data is representative of two independent experiments. **(E)** Representative microscopy images (Left Panel) of sgScramble and sgKdm6a cells treated with 100 µM Etoposide followed by 4 hours of recovery (Bottom Panel) or no recovery (Top Panel) depicting gamma-H2AX foci formation (green-FITC) and nuclei (blue-DAPI). Arrows point to gamma-H2AX foci. Data is representative of two independent experiments. Corresponding box-and-whisker plots (Right Panel) depicting number of gamma-H2AX foci per cell in sgScramble and sgKdm6a cells with 4 hours or no recovery after treatment. (n=30 sgScramble and 44 sgKdm6a Etoposide treated cells (Top) and n=69 sgScramble and 67 sgKdm6a Etoposide treated cells followed by 4 hours of recovery (Bottom)) Two-tailed Student’s t-test was performed. **(F)** Representative histogram (Top Panel) and box-and-whisker plots (Bottom Panels) indicating the MFI of intratumoral gamma-H2AX (Left Panel) and percentage of gamma-H2AX positive tumor cells (Right Panel) in sgScramble and sgKdm6a tumors (n=3 biologically independent samples per group (Left) and n=5 biologically independent samples per group (Right)). Two-tailed Student’s t-test was performed. **(G)** Representative images (Top Panel) displaying comet-like appearance of DNA (vista green) upon single cell alkaline electrophoresis of sgScramble and sgKdm6a cells. Arrows point to comet head and comet tail. Data is indicative of two independent experiments. Box-and-whisker plots (Bottom Panel) representing the Tail DNA %, Tail Moment and Olive Moment of the corresponding comet formed in sgScramble and sgKdm6a cells. (n =246 sgScramble cells and 75 sgKdm6a cells) Two-tailed Student’s t-test was performed. For all the box-and-whisker plots in Figure 3, the centre line marks the median, the edges of the box represent the interquartile (25th–75th) percentile and the whiskers represent minimum-maximum values.

Additionally, we assessed EXO1 protein expression levels in sgScramble and sgKdm6a tumors through immunohistochemical (IHC) staining and imaging of murine tumor sections. This revealed a significant decrease in EXO1 protein expression in sgKdm6a murine tumors (Fig. 3D). To elucidate the functional implications of reduction in EXO1 expression, we conducted several studies to compare genomic integrity between sgScramble and sgKdm6a cells. We first assessed the levels of Gamma(phosphorylated)-H2AX (gamma-H2AX), between sgScramble and sgKdm6a MB49 cells. Gamma-H2AX is a surrogate marker for DNA damage and forms foci at sites of DNA double-strand breaks. The phosphorylation of H2AX at these sites is an early and sensitive indicator of DNA damage response. Confocal microscopy and flow cytometry studies revealed a significantly higher abundance of gamma-H2AX foci in Etoposide (DNA damage inducer)-treated sgKdm6a cells compared to controls (Fig. 3E, Fig. S3E). These findings were validated in tumor samples, where flow cytometry showed significantly higher levels of gamma-H2AX in sgKdm6a tumors (Fig. 3F). Additionally, the COMET assay, which detects DNA damage in individual cells by measuring the migration of fragmented DNA during electrophoresis resulting in a comet-like appearance with a head (intact DNA) and tail (damaged DNA) was used to directly assess DNA breaks (28). The results indicated significantly higher tail DNA percentage, olive tail moment, and tail moment in sgKdm6a cells, demonstrating increased DNA damage (Fig. 3G). Collectively, these data demonstrated that loss of KDM6A impairs DSBR, thereby compromising DNA integrity in bladder cancer cells.

### KDM6A drives glycolysis and lactate production in bladder cancer cells

Previous research has shown that activation of DDR pathways can induce metabolic shifts, including enhanced glycolytic activity to support the Pentose-Phosphate-Pathway and generate nucleotide precursors (29–31). In alignment with reduced expression of genes associated with DDR pathways in KDM6A-Mut patients, we noted attenuated expression of several genes involved in glycolysis and lactate production, including *LDHA, PGAM1*, and *HK2* in KDM6A-Mut patients (Fig. 4A, Fig. S4A). Further, murine ChIP-seq data demonstrated reduced KDM6A binding and concurrent depletion of H3K4me3 enrichment at glycolysis associated genes in sgKdm6a cells (Fig. 4B, C, E, Fig. S4B). Gene set enrichment analysis (GSEA) of H3K4me3 peaks differentially enriched between sgScramble and sgKdm6a cells also showed significant downregulation of the glycolysis pathway with concurrent upregulation of the oxidative phosphorylation (OXPHOS) pathway (Fig. S4C). RNA-seq further corroborated these findings by showing decreased expression of these glycolysis genes in sgKDM6A MB49 cells (Fig. 4D, E). Together, these findings indicate downregulation of the glycolysis pathway in tumor cells following *Kdm6a* deletion.

**Figure 4.**
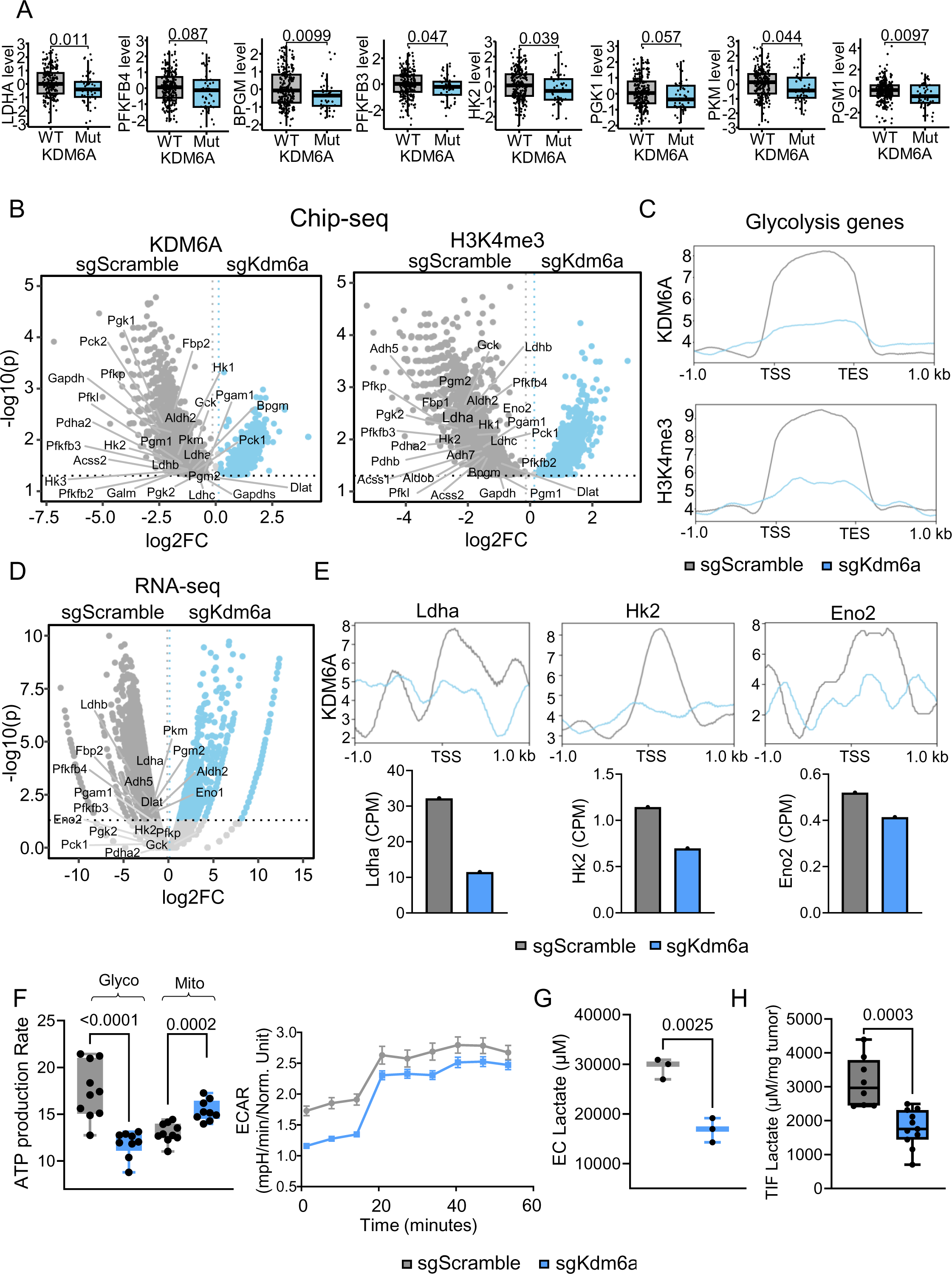
KDM6A drives glycolysis and lactate production in bladder cancer cells. **(A)** Box-and-whisker plots depicting the expression of the indicated genes involved in glycolysis and lactate production in KDM6A-Mut and KDM6A-WT patients, from the IMvigor210 cohort (n=275 patients). Two-tailed Wilcoxon test was performed. **(B)** Volcano plots demonstrating the differential enrichment of KDM6A (Left panel) and H3K4me3 marks (Right panel) in the indicated glycolytic genes between sgScramble and sgKdm6a cell lines, from ChIP–seq data. Volcano plots show the log2 ratio of fold change (log2FC) plotted against the Absolute Confidence, -log10 adjusted p-value (log10(p)). The P-values represent two-tailed probability distribution based on Bayesian model calculated using MAnorm. **(C)** Profile plots representing the probability scores of KDM6A (Top panel) and H3K4me3 (Bottom panel) binding at ±1 kb regions from TSS and TES for genes involved in glycolytic pathway in sgScramble and sgKdm6a cell lines. **(D)** Volcano plot illustrating differentially expressed glycolytic genes between sgScramble and sgKdm6a cell lines from RNA-seq data. Volcano plots depict the log2 ratio of fold change (log2FC) plotted against the Absolute Confidence, -log10 adjusted p-value (log10(p)). P-values were calculated with two tailed exact test under a negative binomial distribution using EdgeR, and adjusted with Benjamini–Hochberg method. **(E)** Profile plots showing the probability scores of KDM6A binding at promoter regions (TSS ±1 kb) of the *Ldha, Hk2* and *Eno2* gene loci in sgScramble and sgKdm6a cell lines (Top panels) and count per million (CPM) bar plots depicting RNA expression levels of the corresponding genes (Bottom panels). **(F)** Box-and-whisker plot (Left panel) representing glycolytic and mitochondrial ATP production rate in sgScramble and sgKdm6a cell lines measured by the Seahorse XF analyzer-based ATP Rate Determination Assay (n=9 biologically independent samples per group). Two-tailed Student’s t test was performed. Line plot (Right panel) comparing Extracellular acidification rate (ECAR) between sgScramble and sgKdm6a cell lines as determined by the Seahorse assay Data represented as mean +/− Standard error of mean (SEM). (n=11 biologically independent samples per group). **(G)** Box-and-whisker plot indicating the Extracellular (EC) L-Lactate concentration in the supernatants of sgScramble and sgKdm6a cell lines, measured following 24 hours of culture (n=3). Two-tailed Student’s t-test was performed. **(H)** Box-and-whisker plot displaying the L-lactate concentrations in the TIF normalized to corresponding tumor weights, obtained from sgScramble and sgKdm6a tumor bearing female mice (n=8 sgScramble and 11 sgKdm6a tumor bearing female mice). Two-tailed Student’s t test was performed. For all the box-and-whisker plots in Figure 4, the centre line marks the median, the edges of the box represent the interquartile (25th–75th) percentile and the whiskers represent minimum-maximum values.

To delineate the functional impact of decreased expression of glycolysis genes in sgKdm6a bladder cancer cells, we compared the metabolic profile of sgScramble and sgKdm6a cell lines using the Seahorse Extracellular Flux Analyzer-based ATP Rate Determination Assay. The data demonstrated reduced glycolytic ATP production rates in sgKdm6a cells compared to controls with concurrent upregulation of mitochondrial OXPHOS (Fig. 4F). Given that downregulation of aerobic glycolysis attenuates lactate production, we compared lactate production by sgScramble and sgKdm6a MB49 cells and observed reduced accumulation of lactate in the culture supernatants of sgKdm6a cells (Fig. 4G). To verify whether these *in-vitro* findings are recapitulated *in-vivo*, we quantified lactate levels in the tumor interstitial fluid (TIF) of sgScramble and sgKdm6a MB49 tumors. We noted significantly lower accumulation of lactate in the sgKdm6a TIF, normalized to tumor weight (Fig. 4H, Fig. S4D), demonstrating that the absence of KDM6A attenuates glycolysis and lactate production in bladder tumors. Importantly, the observed upregulation of the OXPHOS pathway in the cisplatin resistant sgKdm6a cells is in alignment with previous studies associating a switch from glycolysis to OXPHOS pathways with cisplatin resistance. Cumulatively, these findings highlight the role of KDM6A as a major driver of metabolic pathways in bladder cancer and underscore the association between KDM6A deficiency and reduced lactate production by tumor cells.

### Loss of KDM6A-mediated reduction in intratumoral lactate levels reduce Treg abundance and function

Lactate produced by tumor cells has been associated with immune suppression across various tumor types (32). Consequently, we investigated how reduced lactate accumulation in sgKdm6a tumors affects the tumor immune microenvironment and response to anti-PD-1 therapy. Mass cytometry (CyTOF)-based immunophenotyping of sgScramble and sgKdm6a tumors followed by unsupervised clustering and t-SNE analyses of CD45*+* immune cell subsets revealed the presence of distinct T cells, B cells, NK cells, dendritic cells, and immune-suppressive myeloid cell clusters in the TME (Fig. 5A, Fig. S5A). Further, this analysis revealed that sgScramble tumors, which have higher levels of intratumoral lactate, harbor a higher abundance of TGFβ+CD44+LY6C+ immune-suppressive myeloid cells compared to sgKdm6a tumors (Fig. 5B). Further, sgScramble tumors showed a higher abundance of Tregs and increased expression of PD-1 in Tregs compared to sgKdm6a tumors (Fig. 5C, D). Notably, following anti-PD-1 therapy, the Treg population significantly expanded in sgScramble tumors, while this expansion was not observed in sgKdm6a tumors (Fig. 5C). This differential expansion of Tregs resulted in a higher T-effector to Treg ratio in sgKdm6a tumors, indicating a preferential pro-inflammatory shift in the tumor immune microenvironment following anti-PD-1 therapy thus improving responses to ICT (Fig. 5E, F).

**Figure 5.**
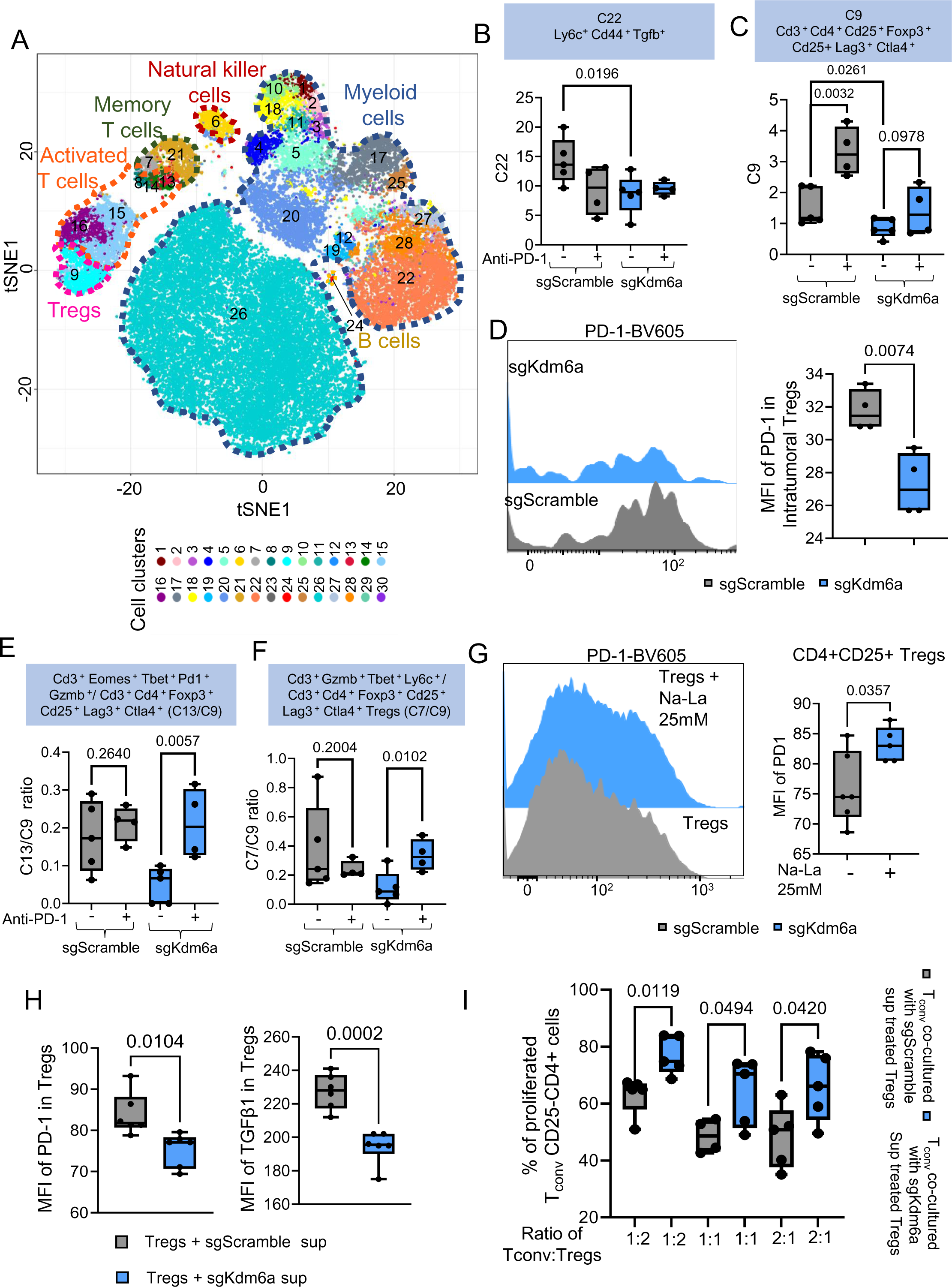
Loss of KDM6A-mediated reduction in intratumoral lactate levels reduce Treg abundance and function. **(A)** t-SNE plot of CyTOF data illustrating the different CD45^+^ immune cell subsets in the Tumor microenvironment (TME) of sgScramble and sgKdm6a tumor bearing female mice treated with and without anti-PD-1. **(B)** Box-and-whisker plot showing relative frequencies of intratumoral Ly6c^+^Cd44^+^Tgfb^+^ Monocytes in female mice bearing sgScramble and sgKdm6a tumors treated with and without anti-PD-1 as assessed by CyTOF. (n=4 female mice for the untreated groups and 5 female mice for the PD-1 treated groups) One-tailed Student’s t-test was performed. **(C)** Box-and-whisker plot displaying relative frequencies of intratumoral Cd3+Cd4+Foxp3+ Cd25+Lag3+Ctla4+ Tregs in sgScramble and sgKdm6a tumor bearing female mice treated with and without anti-PD-1 as determined from CyTOF (n=4 female mice for the untreated groups and 5 female mice for the PD-1 treated groups). One-tailed Student’s t-test was performed. **(D)** Representative histogram (Left Panel) and box-and-whisker plot (Right Panel) representing the MFI of PD-1 in intratumoral Tregs derived from female mice bearing sgScramble and sgKdm6a tumors (n=4 female mice per group). Two-tailed Student’s t test was performed. **(E-F)** Box-and-whisker plots depicting the ratio of intratumoral Cd3^+^Eomes^+^Tbet^+^ Pd1^+^Gzmb^+^ T effector to Cd3^+^Cd4^+^Foxp3^+^Cd25^+^Lag3^+^Ctla4^+^ Tregs (**E**) and the ratio of intratumoral Cd3^+^Gzmb^+^Tbet^+^Ly6c^+^ Memory T cells to Cd3^+^Cd4^+^Foxp3^+^Cd25^+^Lag3^+^Ctla4^+^ Tregs (**F**) in sgScramble and sgKdm6a tumor bearing female mice with and without anti-PD-1 treatment as assessed by CyTOF. (n=4 female mice for the untreated groups and 5 mice for the PD-1 treated groups) One-tailed Student’s t-test was performed. **(G)** Representative histogram (Left Panel) and box-and-whisker plot (Right Panel) demonstrating the MFI for PD-1 in *in-vitro* generated Tregs treated with and without 25 mM Na-La for 48 hours. (n=5 biologically independent samples per group) Two-tailed Student’s t test was performed. **(H)** Box-and-whisker plots indicating the MFI of PD-1 (Left panel) and TGF-β (Right panel) in *in-vitro* generated Tregs treated with the supernatants from sgScramble or sgKdm6a cell lines for 24 hours. (n=6 biologically independent samples per group) Two-tailed Student’s t test was performed. **(I)** Box-and-whisker plot representing the percentage of proliferated, anti-CD3/CD28 Dynabeads stimulated T_conv_ (CD4^+^CD25^−^) cells co-cultured with sgScramble or sgKdm6a supernatant pre-treated Tregs (CD4^+^CD25^+^) at the indicated ratios for 4 days. (n=4 biologically independent samples per group) Two-tailed Student’s t test was performed. For all the box-and-whisker plots in Figure 5, the centre line marks the median, the edges of the box represent the interquartile (25th–75th) percentile and the whiskers represent minimum-maximum values.

Next, to determine the impact of lactate specifically on Tregs, we performed a pHRodo-based lactic acid uptake assay which confirmed that Tregs can uptake exogenous lactate (La). This is in alignment with a previous study that demonstrated that Tregs can utilize exogenous lactate to maintain its functions (33) (Fig. S5B). We also noted that treatment with exogenous sodium lactate (Na-La) increased PD-1 expression in Tregs (Fig. 5G, Fig. S5C). Importantly, when Tregs were treated with supernatants from sgScramble and sgKdm6a cells, those treated with sgKdm6a supernatants had lower PD-1 expression and fewer PD-1+ Tregs linked to the lower lactate levels in the sgKdm6a supernatants (Fig. 5H, Fig. S5D, E). Together, these findings demonstrated that loss of KDM6A decreased glycolysis and lactate production in tumor cells which subsequently reduced PD-1 expression and expansion of PD-1^hi^ intratumoral Tregs following anti-PD1 therapy thus improving the efficacy of anti-PD-1 therapy. In addition to reduction in PD-1 expression, TGFB expression was also notably reduced in sgKdm6a supernatant treated Tregs (Fig. 5H). Production of TGFB is one of the mechanisms through which Tregs exert their immunosuppression (34). Therefore, to determine the impact of extracellular lactate levels on the immune-suppressive function of Tregs, we conducted *in-vitro* T cell suppression assays. We noted decreased proliferation of CD4+CD25-T conventional cells (T_conv_)when co-cultured with sgScramble supernatant-treated Tregs compared to sgKdm6a supernatant-treated Tregs indicating attenuated suppressive potential of the sgKdm6a Tregs (Fig. 5I, S5F). Cumulatively, these findings demonstrate how tumor cell-specific KDM6A regulates the intratumoral metabolic milieu, which in turn alters the tumor immune microenvironment, including PD-1 expression in Tregs and their immune-suppressive function regulating efficacy of anti-PD-1 therapy in bladder cancer.

### Loss of KDM6A in tumor cells impairs lactate mediated histone lactylation and function of regulatory T cells

We and others have demonstrated the role of lactate-derived histone lactylation in the regulation of phenotype and function of immune cell subsets including macrophages and CD8 T cells (35, 36). To determine lactate-mediated epigenetic changes in Tregs, we performed high-performance liquid chromatography (HPLC)–tandem mass spectrometry (MS/MS) analysis of tryptically digested core histones isolated from murine Tregs. We noted that addition of Na-La increased the abundance of H3 histone lysine lactylation demonstrating the role of exogenous lactate in driving lactylation of histone H3 molecules (Fig. S6A). We have previously highlighted the importance of two specific histone H3 lactylation sites-H3K18la and H3K9la in regulating CD8 T cell function and phenotype (36). To understand the relevance of H3K9la and H3K18la in Tregs, we performed flow-cytometry assays to determine the expression of H3K9la and H3K18la in response to exogenous Na-La which demonstrated increased enrichment H3K9la and H3K18la in Tregs following treatment with exogenous Na-la (Fig. 6A, Fig. S6B). Further, to determine the impact of H3K9la and H3K18la on transcriptional regulation in Tregs, we performed ChIP-seq studies followed by ChromHMM analysis. This analysis revealed that chromatin regions marked by H3K9la and H3K18la are also enriched for other transcription initiation marks including H3K4me3, H3K9ac and H3K27ac in Tregs (Fig. S6C). Further, genomic regions enriched with H3K9la and H3K18la marks were identified as transcription start sites (TSS), TSS proximal regions and CpG islands (Fig. S6C). Importantly, treatment with exogenous Na-La resulted in the enrichment of H3K9la and H3K18la marks at the TSS regions indicating a role of exogenous lactate-derived H3K9la and H3K18la in activating gene transcription (Fig. 6B). However, treatment with exogenous lactate did not induce H3K4me3 or H3K27ac enrichment in the promoter regions, indicating selective regulation of H3K9la and H3K18la by exogenous lactate (Fig. 6B). Next, we annotated the genes showing differential enrichment of H3K9la and H3K18la following treatment with exogenous lactate (Fig. 6C). We noted H3K9la and H3K18la enrichment in multiple genes implicated in the regulation of Treg identity and function including *Tgfb, Il10, Helios, Pdcd1, Lag3, Havcr2* and *Irf4* in Na-La treated Tregs (Fig. 6C). ChIP-qPCR and qPCR studies on Na-La treated Tregs also demonstrated higher H3K9la and H3K18la enrichment with concurrent increase in the expression of genes regulating Treg identity and function, including *Foxp3, Pdcd1* and *Tgfb*, corroborating a role of H3K9la and H3K18la in regulating gene expression in Tregs (Fig. 6D, Fig. S6D).

**Figure 6.**
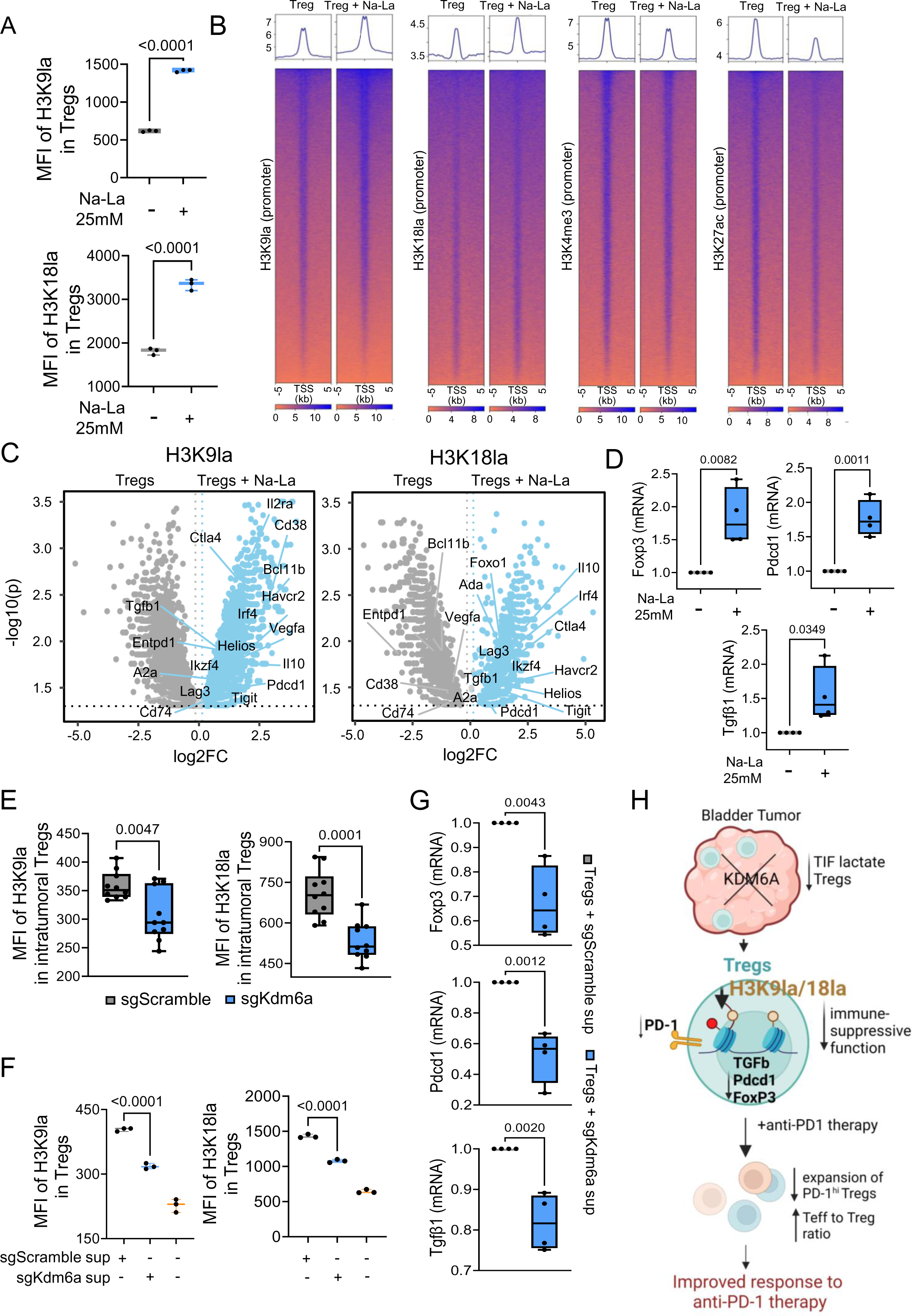
Loss of KDM6A in tumor cells impairs lactate mediated histone lactylation and function of Tregs. **(A)** Box-and-whisker plots depicting the MFI of H3K9la (Top Panel) and H3K18la (Bottom Panel) in *in-vitro* generated Tregs treated with and without 25 mM Na-La for 48 hours. (n=3 biologically independent samples per group) Two-tailed Student’s t test was performed. **(B)** Heatmaps demonstrating the genomic occupancy of the indicated hPTMs at the gene promoter regions in *in-vitro* generated Tregs treated with and without 25 mM Na-La for 48 hours. **(C)** Volcano plots illustrating the differential enrichment of H3K9la (Left Panel) and H3K18la (Right Panel) marks in the indicated genes between *in-vitro* generated untreated Tregs and Tregs treated with 25 mM Na-La for 48 hours, from ChIP-seq study. Volcano plots show the log2 ratio of fold change (log2FC) plotted against the Absolute Confidence, -log10 adjusted p-value (log10(p)). The P-values represent two-tailed probability distribution based on Bayesian model calculated using MAnorm. **(D)** Box-and-whisker plots representing the relative expression of indicated genes (mRNA) in *in-vitro* generated Tregs treated with and without 25 mM Na-La for 48 hours. (n=4 biologically independent samples per group) Two tailed Student’s t-test was performed. **(E)** Box-and-whisker plots showing the MFI of H3K9la (Left Panel) and H3K18la (Right Panel) in intratumoral Tregs derived from sgScramble and sgKdm6a tumor bearing female mice. (n=10 female mice per group) Two-tailed Student’s t test was performed. **(F)** Box-and-whisker plots indicating the MFI of H3K9la (Left Panel) and H3K18la (Right Panel) in *in-vitro* generated Tregs treated with and without culture supernatants from sgScramble and sgKdm6a cell lines for 24 hours. (n=3 biologically independent samples per group). Two-tailed Student’s t test was performed. **(G)** Box-and-whisker plots depicting the relative expression of indicated genes (mRNA) in *in-vitro* generated Tregs treated with sgScramble and sgKdm6a culture supernatants for 24 hours (n=4 biologically independent samples per group). Two tailed Student’s t-test was performed. **(H)** Schematic representation displaying the role of KDM6A in regulating the intratumoral metabolic milieu thus altering the tumor immune microenvironment and driving resistance to anti-PD-1 therapy in bladder cancer. Additionally, the schematic illustrates the role of histone lactylation in modulating Treg phenotype and function within the tumor microenvironment. Schematic was created with BioRender.com. For all the box-and-whisker plots in Figure 6, the centre line marks the median, the edges of the box represent the interquartile (25th–75th) percentile and the whiskers represent minimum-maximum values.

Next, considering the low intratumoral lactate content in sgKdm6a tumors (Fig. S6E), we interrogated the levels of histone lactylation in intratumoral Tregs and noted significantly lower H3K9la and H3K18la in Tregs derived from sgKdm6a tumors (Fig. 6E, Fig. S6F). Further, to directly implicate differences in lactate production by sgScramble and sgKdm6a bladder cancer cells as a major regulator of H3K9la and H3K18la in Tregs, we cultured Tregs in the presence of culture supernatants derived from sgScramble and sgKdm6a cells. Flow cytometry demonstrated significantly lower enrichment of H3K9la and H3K18la in Tregs cultured with KDM6A-deficient supernatants compared to control supernatants, linked to their respective lactate content (Fig. 6F, Fig. S6G, H). Importantly, ChIP-qPCR studies demonstrated attenuated H3K9la and H3K18la enrichment in *Foxp3*, *Tgfb* and *Pdcd1* in Tregs cultured with sgKdm6a cell supernatant (Fig. S6I). Additionally, qPCR studies confirmed reduced expression of these genes, establishing a direct association between extracellular lactate content, H3K9la/H3K18la enrichment, and gene expression including that of *Pdcd1* in Tregs (Fig. 6G). Overall, these findings identified a critical role of the exogenous lactate derived histone PTM-histone lactylation in regulating the expression of critical Treg specific genes, thus regulating their function and response to immunotherapy in sgKdm6a tumors (Fig. 6H).

## Discussion

Therapeutic options for advanced bladder cancer have undergone significant changes in recent years with multiple approvals of anti-PD-1/L-1 therapy, targeted therapies including enfortumab vedotin, sacituzumab govitecan, and combinations of ICT with targeted therapy or chemotherapy (7, 37). However, a major challenge has been identifying biological attributes to select patients for specific therapies and to develop personalized treatment algorithms. Inactivating mutations in *KDM6A,* a commonly mutated gene, is a major driver of bladder cancer initiation and progression. Nonsense, missense, frameshift-indel and splice-site mutations in the *KDM6A* gene which attenuate gene expression are frequent in patients with muscle invasive bladder cancer (38, 39). While *KDM6A* mutations correlate with poor prognosis in bladder cancer, their role in regulating therapeutic outcomes remains unknown. This study investigated the mechanistic underpinnings of how mutations in *KDM6A* impact sensitivity to different therapeutic perturbations. Retrospective analyses of various patient cohorts demonstrated that while mutation in *KDM6A* correlates with improved OS with ICT, it lowers OS with cisplatin-based chemotherapy. Based on these clinical correlations, we adopted a reverse translational strategy to garner mechanistic insights on how the loss of KDM6A impacts tumor cell intrinsic and extrinsic pathways governing response to ICT and cisplatin-based chemotherapy. Pre-clinical studies using CRISPR-Cas9 mediated deletion of *Kdm6a* in murine tumor cells demonstrated that KDM6A is an upstream regulator of multiple tumor cell-intrinsic pathways including DDR and cellular metabolism, which in turn alter the tumor immune microenvironment thus regulating response to ICT and cisplatin-based chemotherapy in bladder cancer.

In this study, we identified KDM6A as a critical regulator of the MMR pathway in bladder cancer. Importantly, we showed that KDM6A directly binds to and regulates the expression of multiple MMR genes. In the absence of KDM6A, the expression of these genes was significantly reduced, thereby disrupting the MMR pathway. Regulation of gene expression is governed by a dynamic interplay of different histone PTMs including H3K27me3 which represses gene transcription and H3K4me3 which creates a permissive chromatin state for transcription initiation (40, 41). KDM6A is known to demethylate H3K27me3 thereby initiating gene transcription. However, recent studies have also shown that knockdown of KDM6A in non-small cell lung cancer cell lines significantly reduces H3K4me3 enrichment, potentially through interactions with the histone lysine methyltransferase KMT2B (42, 43). Consistent with these findings, we have noted that CRISPR-mediated deletion of *Kdm6a* attenuates H3K4me3 enrichment in tumor cells. Overall, our findings demonstrated that loss of *Kdm6a* correlated with decreased H3K4me3 enrichment and attenuated expression of genes associated with the MMR pathway.

We also delineated a critical role of KDM6A in regulating the DSBR machinery including homologous recombination (HR) and Non-homologous end joining (NHEJ) in bladder cancer cells. This is in alignment with previous findings in hematologic malignancies (44). Tumors with defective DSBR machinery rely heavily on PARP-mediated repair for survival which makes them sensitive to PARP inhibition (45, 46). Therefore, future preclinical and clinical studies are warranted to delineate the efficacy of PARP inhibitors either as monotherapy or as combination therapy with ICT in patients with metastatic bladder cancer harboring *KDM6A* mutated tumors which are deficient in DSBR and MMR pathways. We observed that patients with advanced bladder cancer harboring *KDM6A* mutations exhibit decreased expression of MMR and DSBR genes. However, we did not find any correlation between *KDM6A* mutations and mutations in MMR and DSBR genes in these patients. These findings demonstrate that *KDM6A* mutations downregulate critical DNA repair pathways even in the absence of mutations in DSBR and MMR genes. This implicates KDM6A as an independent regulator of the DSBR and MMR pathways in bladder cancer.

Cisplatin based chemotherapy is a major backbone in the treatment of bladder cancer (47). These platinum-based compounds create DNA adducts, causing double-stranded DNA breaks and subsequent tumor cell death. Tumor cells respond by upregulating the DNA repair machinery to resist cisplatin induced killing (48). However, our study demonstrated that sgKdm6a tumors, despite having a deficient DNA damage repair machinery, still resist cisplatin-induced killing. Interestingly, clinical trials have shown that patients with MSI-h tumors, which lack a functional MMR pathway, are also resistant to cisplatin therapy (27). This is because a functional MMR machinery amplifies the impact of cisplatin by attempting futile cycles of repair of cisplatin induced G/T mismatches (25, 26), leading to the generation of additional double-stranded breaks. Therefore, a dysfunctional MMR machinery could potentially explain cisplatin resistance in sgKdm6a tumors. Additionally, cellular metabolism has been linked to cisplatin resistance. Recent studies have shown that a shift from glycolysis to oxidative phosphorylation is associated with increased resistance to cisplatin (49, 50). Consistent with these findings, we observed a similar metabolic switch from glycolysis to oxidative phosphorylation in sgKdm6a bladder cancer cells. Together, the alterations in the MMR and metabolic pathways, could potentially explain the observed resistance to cisplatin-based chemotherapy in sgKdm6a cells and tumors.

Loss of *Kdm6a* attenuated the expression of genes involved in glycolysis, thus suppressing glycolysis-derived lactate generation both *in-vitro* and *in-vivo*. Intratumoral Tregs utilize lactate as a metabolic fuel to support their immune-suppressive functions(33). In alignment with these findings, our CyTOF-based interrogation of the tumor immune microenvironment demonstrated decreased accumulation of Tregs in sgKdm6a tumors which showed reduced intratumoral lactate accumulation. A previous study has reported that lactate plays a role in regulating Treg function through MOESIN lactylation and TGF-beta signaling (51). Additionally, lactate-derived histone lactylation is a histone PTM which has been shown to regulate the phenotype and function of immune cell subsets including macrophages (35). We have previously delineated the role of endogenous glycolysis-derived histone lactylation in regulating the phenotype and function of different CD8 T cell subsets (36). In this study, we demonstrated that tumor-derived lactate regulates histone lactylation in Tregs. A series of experiments demonstrated the enrichment of H3K9la and H3K18la in key genes including *Foxp3*, *Tgfb*, and *Pdcd1*, involved in regulating Treg phenotype and function. Importantly, loss of *Kdm6a*-mediated reduction in intratumoral lactate resulted in attenuation of H3K9la and H3K18la in intratumoral Tregs with concurrent reduction in the expression of T-reg specific gene such as *Foxp3* and *Tgfb*.

Further, we noted reduced abundance of intratumoral PD-1^hi^ T regs in mice bearing sgKDM6A tumors. A previous study indicated that lactate regulates PD-1 expression in Tregs (52). Therefore, reduced abundance of intratumoral lactate in mice bearing sgKDM6A tumors possibly attenuated histone lactylation-mediated expression of PD-1 in Tregs. The expansion of PD-1^hi^ Tregs has been linked to resistance to anti-PD-1 therapy. Failure to expand the PD-1^hi^ T regs in mice bearing sgKDM6A tumors increased the ratio of cytotoxic CD8 T cells to Tregs with improved response to anti-PD-1 therapy highlighting the potential role of lactate mediated histone lactylation in regulating the response. Together, these findings uncovered a previously unrecognized role of histone lactylation in modulating Treg phenotype and function within the TME. Additionally, these findings delineated the role of KDM6A in regulating the intratumoral metabolic milieu thus altering the tumor immune epigenome and driving response to ICT in bladder cancer.

Overall, our study provides a mechanistic framework to utilize *KDM6A* mutation for patient stratification and development of personalized treatment algorithms.

## Methods

### Cell lines (CRISPR-Cas9 mediated deletion of *Kdm6a*)

The parental MB49 cell line was obtained from Sigma-Aldrich and authenticated by American Type Culture Collection (ATCC) using Short Tandem Repeat (STR) profiling. CRISPR-Cas9 technology was used to excise out the *Kdm6a* gene from the parental MB49 cell line to generate the sgKdm6a cell line by the MDACC Functional Genomics Core. Briefly, single guide RNA (sequence: TAGCATTATCTGCATACCAG) was subcloned into LentiCRISPRv2 vector and subsequently transfected into MB49 cells using the JetPrime transfection reagent. 48-72 hours 2 days after transfection, 2ug/ml of puromycin was used to remove non-transfected cells. The sgKdm6a cell line was screened by western blots and genomic DNA sequencing. The control cell line, sgScramble was generated by first transfecting LentiCRISPRv2 only with pMD2.G and psPAX2-D64V vectors into 293T cells. 48 hours after transfection, the virus-containing medium was collected and used for transduction with MB49 parental cells. 72 hours following transduction, 2ug/ml of puromycin was used to remove non-transfected cells.

### Mice

This study used age-matched 5–7-weeks old female C57BL/6 mice for experiments. Mice were housed and sustained in pathogen-free conditions, 20-25°C, 30-70% humidity and on a 12-hour light/12-hour dark cycle at the Animal Resource Center, UTMDACC. The UTMDACC Institutional Animal Care and Use Committee approved all animal protocols.

### Patients and samples

The publicly available dataset for IMvigor210, a phase 2 study evaluating atezolizumab in advanced bladder cancer (NCT02108652, NCT02951767), was accessed through the IMvigor210CoreBiologies R package (Mariathasan et al., 2018). The impact of *KDM6A* mutation status on OS, TMB, and the expression of glycolysis-related genes among advanced bladder cancer receiving ICT (n = 275) was studied. Additionally, the OS data for advanced bladder cancer patients (with Oncotree code “BLCA”, n =148) treated with ICT was retrieved from cBioPortal (Samstein et al., 2019). This study data included patients who received at least one dose of ICT (atezolizumab, avelumab, durvalumab, ipilimumab, nivolumab, pembrolizumab, or tremelimumab). We compared expression levels of gene involved in MMR, DSBR and glycolytic pathways from between *KDM6A*-Mut and *KDM6A*-WT for advanced bladder cancer patients treated with immunotherapy utilizing transcriptomic data the retrieved from (Damrauer et al., 2022).

The clinical and genomic data for patients with resectable and advanced bladder cancer receiving cisplatin-based chemotherapy were accessed from two independent studies, blca_msk_tcga_2020 and paired_bladder_2022 (Pietzak et al., 2019, Clinton et al., 2022) retrieved from cBioPortal.

Survival outcomes were analyzed using Cox proportional hazards (PH) regression models, with PH assumptions validated through scaled Schoenfeld residuals. The magnitude of associations was quantified using hazard ratios (HRs). Kaplan-Meier plots were generated to visually depict the relationship between KDM6A mutational status and OS using “ggsurvplot” function in R.

TMB for patients with advanced bladder cancer was retrieved from The Cancer Genome Atlas (TCGA-BLCA, n = 136) and mutational burden was compared between patients with and without KDM6A mutations (4). Somatic variants summary for all bladder cancer patients in the TCGA-BLCA (n= 412) were downloaded in Mutation Annotation Format (maf) using TCGAbiolinks R package. The co-occurrence or mutual exclusivity of *KDM6A* mutations with mutations in MMR and DSBR-related genes was computed using the “somaticInteractions” function in maftools v2.10.05. All statistical analyses were conducted using R v4.1.2 and GraphPad Prism 9.0.

### Western Blot

sgScramble and sgKdm6a cells lines in culture were trypsinized and thoroughly resuspended in RIPA buffer supplemented with protease inhibitor by vortexing every 5 minutes for 25 minutes on ice. Afterward, samples were centrifuged at 12,000 rpm to pellet any debris and the protein in the collected supernatant was quantified using the Pierce BCA Protein Assay kit according to the manufacturer’s protocol. Following protein quantification, 15 μg of protein were loaded into a PROTEAN TGX gel and underwent electrophoresis for 1 hour. After gel electrophoresis, the proteins were transferred to an Immobilon membrane for 1 hour. Then, the membrane with the transferred protein was blocked for 1 hour in EveryBlot Blocking buffer, then incubated overnight at 4°C in 1:2000 of anti-KDM6A primary antibody diluted in EveryBlot Blocking Buffer. The following morning, the membrane was washed 5 times in PBST before being incubated with 1:2000 of anti-Rabbit IgG HRP linked secondary antibody diluted in EveryBlot Blocking Buffer for 1 hour. Lastly, the membrane was washed in PBST six times followed by chemiluminescent imaging in a BioRad ChemiDoc imaging instrument.

### *In-vivo* tumor model and tumor processing

sgScramble and sgKdm6a cells were collected from culture via trypsinization during logarithmic phase of growth and then washed with PBS. C57BL/6 mice were ijected subcutaneously in the right flank with 2 × 10^5^ sgScramble or sgKdm6a cells resuspended in 100 μl of PBS. To assess tumor growth, tumors were measured by Vernier Calipers. On Day 10 or 11 post tumor implantation, mice were sacrificed, and the tumors were isolated and weighed.

Collected tumors were transferred to Eppendorf tubes containing FBS-free media supplemented with 0.66 mg/ml of Liberase and 20 mg/ml of DNAse. Next, tumors were chopped using scissors,and the sample tubes were transferred to a thermal shaker for tumor digestion at 37°C at 1,000 rpm for 30 minutes. Following tumor digestion, a single-cell suspension was obtained by passing tumors through a 70 μm strainer atop a 50 ml conical tube using the back of a 1ml syringe. The single-cell suspension was subsequently washed in PBS at 200 x g for 5 minutes at 4°C, the cell pellet was subjected to RBC lysis by resuspending in ACK Lysis Buffer and then washed in PBS before proceeding to downstream assays. For some experiments, the cell pellet was resuspended in freezing media comprising of 90% FBS and 10% DMSO before storage at −80°C for downstream use.

### Anti-PD-1 *in-vivo* treatment regimen

Tumor-bearing mice were injected intraperitoneally with 200 μg, 100 μg, and 100 μg of anti-PD1 diluted in 100 ul PBS on Days 3, 6 and 9 post-tumor implantation. Tumors were measured using a Vernier Caliper on Days 6 and 9 to assess tumor growth. The tumors were isolated, weighed, and subjected to tumor processing on Day 10 or 11 post-tumor implantation as described previously.

### *In-vivo* cisplatin-based chemotherapy regimen

Tumor-bearing mice were injected intraperitoneally with 2.5 mg/kg of gemcitabine plus 6 mg/kg of cisplatin on Days 1, 4 and 7 post-tumor implantation. Tumor volumes were measured by Vernier Caliper on Days 4, 6 and 8. Tumors were isolated, weighed and subjected to tumor processing on Day 10 or 11 post-tumor implantation as described previously.

### Murine splenocyte isolation and *in-vitro* Treg generation

Spleens were harvested from female C57BL/6 mice, rinsed with cold PBS, mushed, and filtered using a 70 µm strainer. The resulting single-cell suspension was washed once with cold PBS by centrifugation and treated with ACK Lysis Buffer to lyse the RBCs. The cell counts were obtained using an automated cell counter before proceeding to Magnetic Assisted Cell Sorting (MACS) of naïve CD4 T cells.

Naïve CD4 T cells were magnetically isolated using the Murine Naïve CD4 T Cell Isolation Kit, as per manufacturer’s guidelines. Briefly, splenocytes were resuspended in MACS buffer (1X PBS supplemented with 0.5% BSA and 2mM EDTA) on ice. Naïve CD4 Biotin Antibody Cocktail was added, mixed thoroughly, and incubated at 4°C for 15 minutes. Next, Anti-biotin Microbeads were added to the samples, mixed thoroughly, and incubated on ice for an additional 15 minutes. CD44 Microbeads were then added, followed by a final incubation for 20 minutes on ice. The samples were subsequently washed by centrifugation at 200 X g for 5 minutes at 4°C in surplus MACS Buffer. The resulting cell pellets were resuspended in MACS buffer, passed through LS columns with the naïve CD4 T cells been collected in the flow-through.

Subsequently, to generate Tregs, these naïve CD4 T cells (1-1.5 × 10^6^ cells per well) were stimulated with 3 μg/ml of anti-CD3, 2 μg/ml of anti-CD28, 10 ng/ml of murine recombinant IL-2, and 10 ng/ml of recombinant mouse TGF-β in complete RPMI media and plated in 24-well plates. The cells were then incubated for 5 days at 37°C with 5% CO_2_. On day 5, the percentage of Foxp3^+^ Tregs generated *in-vitro* was examined using flow cytometry as described below.

### ChIP Sequencing

ChIP assays were performed on sgScramble and sgKdm6a cell lines, as well as *in-vitro* generated Tregs, treated with or without 25 mM Na-La for 48 hours, using the MAGnify Chromatin Immunoprecipitation System kit as per the manufacturer’s instructions. Briefly, cells were incubated with 37% formaldehyde for 10 minutes at room temperature to crosslink the chromatin. Then to stop the crosslinking reaction, the samples were treated with 1.5 M glycine and incubated for 5 minutes at room temperature. Next, the samples were washed thrice with ice-cold PBS through centrifugation at 200Xg at 4°C for 10 minutes. The resulting cell pellets were then resuspended in lysis buffer supplemented with protease inhibitors and subjected to sonication to fragment the DNA into 150-300 kb fragments. For every immunoprecipitation reaction, 10 μg of specific antibodies as detailed in the figure legends, were first coupled with Protein A/G Dynabeads and then added to the sheared samples and incubated overnight at 4°C.The subsequent day, unbound antibodies were removed by washing thrice with Immunoprecipitation (IP) Buffer 1 and twice with IP Buffer 2. The crosslinking was then reversed through heat treatment, using the Reverse crosslinking buffer supplemented with Proteinase K. Next, the target DNA was purified using the DNA purification magnetic beads resuspended in DNA purification buffer and incubated at room temperature for 30 minutes. The bead-bound DNA was then washed twice with DNA wash buffer to remove any residual contaminants. Finally, the purified target DNA was eluted out by incubating the bead-bound DNA with DNA elution buffer at 55°C for 20 minutes in a thermocycler. The immunoprecipitated DNA was then quantified with the dsDNA HS Assay kit using the Qubit Flex Fluorometer. 10 μg of DNA from each condition was sent to the MDACC Advanced Technology Genomics Core (ATGC) for library preparation and sequencing which was carried out using the Illumina NextSeq500 instrument.

### ChIP Sequencing data analysis

The quality of CHIP-seq FASTQ sequences generated as described above, were assessed using FastQC v0.11.9, followed by mapping by bowtie2 v2.3.5.1 (53)(Langmead and Salzberg, 2012) with mouse reference genome mm10. The bam files obtained from mapping were further processed using SAMBLASTER v0.1.26 (54) and SAMTOOLS v1.13 (55), for duplicate removal, sorting and indexing. Further, SAMBAMBA v0.6.6 (56) was used to normalize the bam files per read counts by performing random sampling. The ChIP-seq signal enrichment over “Input control” background was identified using Model based analysis of ChIP-seq (MACS)2 v2.2.8 (57). The peaks with significant p values (< 0.05) were considered for further annotation with using CHIPseeker v1.30.3 (58) and clusterProfiler (59) packages in R v4.3.2. The quantitative pairwise comparisons of different datasets were performed using Manorm v1.1.4 (60) using both peak (bed) and read (bam) coordinates for respective sample. The significant enrichment of target genes was estimated of the basis of M value which describes the log2 fold change and plotted with ggplot2. Differential pathways enriched among datasets were identified using GSEA v4.2.3 (61). The bigwig files were generated from aligned bam reads with “bamCoverage” function of deepTools v3.5.1 (62) and profile plots for specific genes were plotted using the computeMatrix and plotProfile programs of deepTools. The genome wide annotation of aligned reads was performed using ChromHMM v1.24 (63) which applies multivariate hidden Markov model (HMM) to assign states by modeling combinatorial presence and absence of each mark. The reads were subjected to “BinarizeBam” followed by “LearnModel” using mm10 assembly study enrichment and functional annotation of each mark in 7-state model.

### RNA sequencing

sgScramble and sgKdm6a cells were collected from culture at the logarithmic phase of growth by trypsinization, washed once in PBS, and then snap-frozen in liquid nitrogen preceding total RNA extraction performed by the MDACC Biospecimen Extraction Core Facility. A minimum of 120 ng of extracted RNA with an RNA Integrity Number value >7 was sent to the MDACC ATGC for stranded, paired-end mRNA sequencing in a NextSeq500 instrument.

### RNA sequencing data analysis

The paired-end raw FASTQ sequences were subjected to quality control with FastQC v0.11.9, followed by adapter removal using Trimmomatic v0.39(Bolger et al., 2014). The trimmed fastq reads were mapped to mm10 reference genome using STAR v2.7.3a(Dobin et al., 2013) and resulting bam reads were sorted and indexed with SAMTOOLS v1.13. The gene counts for each sample were estimated with HTseq v2.0.2(Putri et al., 2022) and the counts were converted to “counts per million (CPM)” for differential analysis between samples with EdgeR v3.36.0(Robinson et al., 2010). The regulatory potential scores of Chipseq peaks of different antibodies for expressed genes were calculated by integrating Chipseq experiments and RNA-seq differential expression data using BETA v1.0.7(Wang et al., 2013). The heatmaps were plotted with ggplot2 using R v4.3.2.

### Isolation of CD4^+^CD25^+^ Tregs and CD4^+^CD25^−^ T_conv_ cells

Murine CD4^+^CD25^+^ Tregs, and CD4^+^CD25^−^ T_conv_ cells were magnetically isolated using murine CD4^+^CD25^+^ Regulatory T Cell Isolation Kit, according to the manufacturer’s protocol. Briefly, cells were resuspended in MACS buffer and CD4^+^CD25^+^ Regulatory T Cell Biotin-Antibody Cocktail was added and incubated on ice for 15 minutes. Next, the Anti-Biotin Microbeads were added, incubated at 4°C for 15 minutes, following which the samples were washed in surplus MACS buffer by centrifugation. The resulting cell pellet were passed through the LD columns, flow through was collected and centrifuged at 200 X g for 5 minutes at 4°C. The cell pellet was then resuspended in MACS buffer, CD25-PE antibody was added, mixed well, and incubated for another 15 minutes on ice. Subsequently, the samples were incubated for an additional 15 minutes with Anti-PE Microbeads and washed in excess MACS buffer by centrifugation. The pellets were resuspended and run through the MS Columns. CD4^+^CD25^−^ T_conv_ cells were collected in the flow-through, while CD4^+^CD25^+^ Tregs were eluted from the column using a plunger.

### Flow cytometry

Flow cytometry was conducted on several experimental groups, including: sgScramble and sgKdm6a cell lines, *in-vivo* generated tumors derived from these cell lines, *in-vitro* generated Tregs treated with and without 25 mM Na-La for 48 hours and, *in-vitro* generated and CD4^+^CD25^+^ sorted Tregs treated with the supernatants from sgScramble and sgKdm6a cell lines for 24 hours.2-3 × 10^6^ of each of the above-mentioned cell types were briefly rinsed in FACS buffer (PBS supplemented with 5% FBS) by centrifugation at 200 x g at 4°C for 5 minutes. After washing, the cells were incubated on ice for 15 minutes in a blocking buffer containing murine, bovine, hamster, rat, and rabbit serum along with 25 μg/mL 2.4G2 (Fc block) antibody in PBS. The cells were then stained with the surface antibody cocktail for 30 minutes at 4°C. After, two subsequent washes with FACS buffer, the cells were fixed and permeabilized on ice for 45 minutes. Following another wash with 1X permeabilization buffer, the cells were incubated in the intracellular antibody cocktail as mentioned in for 20 minutes at room temperature. Finally, the samples were washed with FACS buffer, fixed with 1% paraformaldehyde. The samples were then acquired using a BD LSRFortessa flow cytometer and FlowJo v10.9 software was used for analysis of the flow cytometry data.

### Microsatellite Instability detection

To detect microsatellite instability (MSI) in sgScramble and sgKdm6a tumor cells, microsatellite marker mBAT-64 was PCR amplified from the genome, and the size of the marker was analyzed by fluorescent fragment length analysis (FFLA). Briefly, sgScramble and sgKdm6a tumors were processed as described previously to obtain sgScramble and sgKdm6a tumor cells, 2 ×10^6^ cells resuspended in PBS were transferred to Eppendorf tubes and centrifuged at 2000 rpm for 5 minutes at 4°C. Genomic DNA was extracted from tumor cells using QIAamp DNA Micro Kit, according to the manufacturer’s instructions. For cell lysis, the cell pellet was resuspended in buffer ATL, proteinase K and buffer AL was added and pulse-vortexed for 15 seconds, followed by 10 minutes incubation at 56°C in a shaker. After incubation, 100% ethanol was added, pulse-vortexed for 15 seconds and incubated at room temperature for 3 minutes. The lysates were then transferred to QIAamp MinElute columns and centrifuged at 6000 x g for 1 minute at room temperature, followed by washing with buffer AW1 and AW2. Then, the columns were centrifuged at 20,000 x g to remove excess buffer, nuclease-free water was added incubated for 5 minutes at room temperature and centrifuged at 20,000 x g to elute the genomic DNA. The concentration of genomic DNA was measured with a Qubit Flex Fluorometer using the dsDNA HS Assay kit according to the manufacturer’s instruction. The microsatellite marker *mBAT-64* was PCR amplified from 20ng of genomic DNA with a 5’ PET-tagged forward primer and reverse primer, using the Type-it Microsatellite PCR Kit as per the manufacturer’s instructions. PCR condition was as follows: initial activation at 95°C for 5 minutes; followed by 27 cycles of 95°C for 30 seconds, 60°C for 90 seconds and 72°C for 30 seconds; then a final extension step at 60°C for 30 minutes. The size and distribution of *mBAT-64* PCR amplicons was determined by capillary electrophoresis performed on a 3730 Genetic Analyzer instrument followed by analysis of data using the Peak Scanner software. The PET-tagged *mBAT-64* amplicons were sent to MDACC ATGC for FFLA. The amplicons were mixed with formamide and Liz 500 size standard in a 96-well PCR plate, loaded on the mentioned instrument and run for 45 minutes. The results(.fcs files) were imported to Peak Scanner software, size standard and analysis method were set to GS500 LIZ and Sizing Default PP, respectively and analyzed. The *mBAT-*64 modal peaks in the electropherogram were identified between 110-140 bp and the modal peak position in base pairs were compared between sgScramble and sgKdm6a tumor samples.

### Annexin V based cytotoxicity assay

To study cisplatin-induced cytotoxicity in sgScramble and sgKdm6a cells, the cells were treated with or without 5 µM cisplatin for 48 hours, following which the cells were harvested and stained with Annexin V-Pacific Blue diluted in 1X Annexin Binding Buffer for 15 minutes at room temperature. The stained cells were then washed with 1X Annexin Binding Buffer and acquired immediately on the BD LSR Fortessa flow cytometer and analyzed using FlowJo v10.9 software.

### Spheroid generation assay

To compare the spheroid formation ability of sgScramble and sgKdm6a cells in presence of cisplatin, cultured cells were allowed to form spheroids on top of agarose in presence of 10 µM cisplatin. Briefly, 100 µl of 1.5% molten agarose was transferred to each well of a 96-well flat-bottom plate and allowed to solidify. Then 2 × 10^4^ cells resuspended in 200µl of complete DMEM media containing 10µM cisplatin were transferred to the top of the agarose. To allow spheroid formation, the plates were centrifuged at 220 x g for 10 minutes and incubated for 48 hours at 37°C. After 48 hours, the formed spheroids were imaged in a EVOS FL imaging system under a 4X objective. The ImageJ software was used to measure the diameter of the spheroids using the scale in the microscopy images as reference.

### Wound-healing assay

To compare the migration ability of sgScramble and sgKdm6a cells in presence of cisplatin, wound-healing assay was performed. 2 ×10^5^ sgScramble and sgKdm6a cells were seeded into each well of a 24-well plate and incubated until cells reached confluency. Using a sterile 200 µl pipette tip a scratch was made through the middle of the well and imaged at 0 hours in a EVOS FL imaging system under a 4X objective. The cells were then treated with or without 5 µM cisplatin followed by 24 hours incubation and imaged at 24 hours as described previously. The images were analyzed in ImageJ Software using the Wound healing size tool plugin to determine the wound closure percentage. The scratch area was determined at 0 hours and 24 hours by using the following parameters of plugin: Variance window radius, threshold value and percentage of saturated pixel set to 10, 100 and 0.001 respectively, and set scale global and the scratch is diagonal was enabled. The percentage of wound closure was determined by calculating the percentage decrease in the scratch area between 0 and 24 hours.

### Immunohistochemistry (IHC)

IHC staining was performed on formalin-fixed, paraffin-embedded tissue sections of sgScramble and sgKdm6a tumors to detect EXO1 levels. sgScramble and sgKdm6a tumors were isolated as described previously, fixed in 10% formalin for 4 hours and sent to MDACC Research Histology Core Laboratory (RHCL) for sectioning and staining. Fixed tumors were embedded in paraffin and sectioned at 4-micron thickness. Next, sections were deparaffinized, hydrated and antigen retrieved with BOND Epitope Retrieval Solution 1 followed by washing with BOND wash buffer. For staining BOND Polymer Refine Detection kit was used, sections were blocked with hydrogen peroxide for 15 minutes, stained with rabbit anti-mouse EXO1 antibody at a 1:200 dilution for 30 minutes, followed by staining with anti-rabbit Poly-HRP-IgG secondary antibody for 15 minutes. Slides were visualized using 3′-3-Diaminobenzidine substrate as a chromogen, followed by hematoxylin counterstaining for 3 minutes. Slides were scanned into Aperio eSlide manager for observation.

### Gamma-H2AX assay of sgScramble and sgKdm6a cells

To detect dsDNA breaks in sgScramble and sgKdm6a cell lines, presence of gamma-H2AX foci were detected using flow cytometry and microscopy-based studies.

For flow cytometry experiments, sgScramble and sgKdm6a cells were seeded at 2 × 10^5^ cells per well in a 24-well plate and incubated overnight at 37°C. The following day, cells were treated with 100µM Etoposide for 1 hour followed by 4 hours recovery or no recovery. 3 × 10^6^ cells resuspended in PBS were transferred to a 96-well round bottom plate, centrifuged at 2000 rpm for 5 minutes at 4°C and resuspended in fixation and permeabilization buffer for 40 minutes on ice. Following fixation and permeabilization, the cells were centrifuged at 2500 rpm for 5 minutes at 4°C and the cell pellet was subsequently stained with Anti-Phospho-Histone H2A.X (Ser 139) primary antibody for 20 minutes at room temperature. After incubation, cells were washed with PBS by centrifugation at 2500 rpm for 5 minutes at 4°C and subsequently stained with anti-mouse FITC-conjugated secondary antibody for 15 minutes in dark at room temperature. Then, the cells were washed with PBS and fixed with 1% paraformaldehyde. The samples were acquired using a BD LSR Fortessa flow cytometer and analyzed using FlowJo v10.9 to determine the percentage of gamma-H2AX positive cells.

For microscopy experiment, 1 × 10^6^ sgScramble and sgKdm6a cells were seeded on poly-D-lysine coated coverslips and incubated overnight at 37°C. The following day, cells were treated as described previously, then fixed with 3.7% paraformaldehyde for 20 minutes at room temperature, permeabilized with 1% Triton X-100 for 10 minutes at room temperature and subsequently blocked for 30 minutes with blocking buffer (1% BSA in PBS) at room temperature. After blocking, the cells were incubated with the H2AX primary antibody as mentioned previously for 90 minutes at room temperature, washed with PBST (0.05% Tween-20 in PBS) and then incubated with anti-mouse secondary antibody in the dark for 1 hour at room temperature. Then, the cells were washed with PBST followed by DAPI staining in the dark for 15 minutes. To remove excess dye cells were washed with PBS and cover slips were mounted on to slides using mounting media. The slides were imaged using Leica SP8 confocal system at the MDACC Advanced Microscopy Core (AMC). The image files (.lif files) were imported to ImageJ software using the Bio-Formats plugin, gamma corrected to a value of 0.35 for further analysis and the number of distinctly visible gamma-H2AX foci were counted manually for each cell.

### Gamma-H2AX assay of *in-vivo* tumor cells

sgScramble and sgKdm6a tumors were processed as described previously to obtain sgScramble and sgKdm6a tumor cells. 3 × 10^6^ sgScramble and sgKdm6a tumor cells were transferred to a 96-well round bottom plate, centrifuged at 2000 rpm for 5 minutes at 4°C and the cell pellet was stained with PB conjugated anti-CD45^+^ antibody followed by 15 minutes incubation at room temperature. The cells were then washed with PBS followed by fixation and permeabilization and staining with H2AX primary antibody and anti-mouse secondary antibody as described previously. The cells were then washed with PBS and incubated with PE conjugated tumor cell marker anti-E-Cadherin antibody for 15 minutes at room temperature. The cells were then washed, fixed, acquired and analyzed as described previously to determine the MFI of gamma-H2AX in tumor cells and percentage of gamma-H2AX positive tumor cells.

### Comet assay

To detect DNA damage in sgScramble and sgKdm6a cell lines, comet assay kit was used for single cell gel electrophoresis and DNA staining according to the manufacturer’s protocol. Cultured cells were harvested, resuspended in 1X cold PBS at 5 × 10^5^ cells/ml and mixed with molten comet agarose at 37° C in the ratio of 1:10 (v/v). A base layer of agarose was prepared on comet slides and allowed to solidify, then the cell-agarose mixture was transferred to the top of the base layer. Following solidification of the cell layer, alkaline lysis of cells was performed by immersing the slides in pre-chilled lysis buffer for 45 minutes at 4°C in the dark, followed by a 45-minute immersion in pre-chilled alkaline solution. Then single cell alkaline electrophoresis was performed by placing the slides in an electrophoresis chamber containing cold alkaline electrophoresis solution, at 1 volt/cm for 25 minutes with a current setting of 310 mA. The slides were then washed three times with cold autoclaved water, followed by cold 70% ethanol for 5 minutes. Slides were allowed to air dry at room temperature and stained with the Vista green DNA dye for 15 minutes in dark at room temperature. Excess dye was removed, and slides were imaged using Leica SP8 confocal system at the MDACC Advanced Microscopy Core. The image files (.lif files) were imported to ImageJ software as described previously, and gamma corrected to a value of 0.35 for further analysis. Using the OpenComet plugin, profile analysis was done with background correction off for comet finding and head finding. The output of the analysis provided the corresponding tail DNA percentage, tail moment and olive moment of the comets.

### Seahorse assay (ATP Real-time Rate Assay)

To compare the ATP production rates from glycolytic and mitochondrial pathways between sgScramble and sgKdm6a cell lines, the Seahorse XF Real-Time ATP rate assay was performed following the manufacturer’s instructions by the MDACC Metabolomics Core. The cells were seeded overnight on the XF-96 well-plate. The assay was performed using an XF-96 Analyzer which measured the extracellular acidification rate (ECAR) under basal conditions as well as after sequential injections of 1.5 μM oligomycin and 0.5 μM Rotenone/ Antimycin A. Real-time rates of ATP production were recorded and assessed using the Wave software.

### Extracellular lactate measurement assay

To distinguish the concentration of L-Lactate produced by sgScramble and sgKdm6a cell lines, culture supernatants were collected from these cell lines following 24 hours of culture.

To compare the intratumoral concentration of L-Lactate in sgScramble and sgKdm6a tumors, tumor interstitial fluid (TIF) was collected by resting tumors on a 40 μm nylon filter atop a 50 ml conical tube, then centrifuging the sample tubes at 106 x g for 10 minutes at 4°C. The collected TIF samples were transferred to separate Eppendorf tubes, snap-frozen in liquid nitrogen, and then stored at −80°C before assaying.

TIF and culture supernatant samples were processed for extracellular lactate quantification using the L-Lactate Assay Kit. Briefly, samples underwent deproteination using cold 0.5M MPA on ice for 5 minutes. After deproteination, samples were centrifuged at 10,000 x g for 5 minutes at 4°C. Next, the supernatant was collected, transferred to a new tube, then subjected to neutralization using potassium carbonate and centrifuged at 10,000 x g for 5 minutes at 4°C, after which the supernatant was collected. The assay plating was conducted following the manufacturer’s instructions, and the absorbance was read using a BioTek Synergy H1 microplate reader at 535 nm.

### Mass cytometry (CyTOF) for immune profiling of tumor microenvironment

Tumor samples processed and stored as previously described were revived by adding samples drop by drop to complete DMEM media. Then, 3 × 10^6^ cells were collected from each sample and washed with PBS, then resuspended in 194Pt monoisotopic cisplatin at a final concentration of 5 μM in FACS buffer and incubated for 3 minutes at room temperature. Next, samples were washed three times with FACS buffer followed by fixation and permeabilization for 15 minutes at room temperature using Maxpar barcode fixation and permeabilization buffer. Then, samples were labeled using palladium barcoding, according to the manufacturer’s protocol, and incubated for 30 minutes at room temperature. After labeling, samples were washed three times in FACS buffer and surface staining was performed by resuspending samples in the antibody mixture prepared in blocking buffer at 4°C for 30 minutes. After surface staining, samples were washed twice with FACS buffer and then subjected to fixation and permeabilization for 1 hour at 4°C. Next, samples were washed twice in permeabilization buffer and resuspended in the intracellular antibody mixture prepared in permeabilization buffer for 30 minutes at 4°C. After washing with FACS three times, samples were fixed using 1.6% paraformaldehyde in PBS supplemented with 100 nM iridium nucleic acid intercalator and left overnight at 4°C. The following day, samples were washed with PBS three times and resuspended in nuclease-free water before acquisition using a Helios mass cytometer. We analyzed the CyTOF data using Premessa, FlowCore, and FlowSOM packages as described in a previously published protocol.

### pHrodo red assay for lactic-acid uptake

Tregs were collected from culture and 1 × 10^6^ cells were plated per well in a 96 well plate, then washed with 20 mM HEPES in PBS. After washing, cells were resuspended in the pHrodo staining solution prepared according to manufacturer’s instructions in 20 mM HEPES in PBS, and incubated at 37°C for 30 minutes. Following staining, cells were washed twice in 20 mM HEPES in PBS. Following the washes, lactic acid was added to each sample well at final concentrations of 5 mM and 15 mM and then incubated at 37°C for 30 minutes, following which samples were acquired on a BD LSR Fortessa flow cytometer and analyzed using FlowJo v10.9.

### *In-vitro* suppression assay

Tregs were generated *in-vitro*, magnetically sorted for CD4^+^CD25^+^ Tregs as described previously and then treated with the supernatant of sgScramble or sgKdm6a cell lines for 36 hours. For T_conv_ cells, single cell suspensions of murine splenocytes were obtained and magnetically isolated for CD4^+^CD25^−^ T_conv_ cells as previously outlined.

T_conv_ and Tregs were labelled with Cell Trace Violet (CTV) and Far Red (FR) respectively according to the manufacturer’s protocol. T_conv_ (5 × 10^4^) cells were co-cultured with Tregs (5 × 10^4^) at varying ratios (1:1,2:1 and 2:1, T_conv_:Treg ratio) and stimulated using anti-mouse CD3/CD28 Dynabeads and 10 ng/ml of murine recombinant IL-2 per the manufacturer’s guidelines. The cells were co-cultured in 96 well round bottom plates with 200 μl RPMI for 4 days at 37°C and 5% CO_2_.

On Day 4, cells were acquired using the BD LSR Fortessa. FlowJo v10.9 software was employed to analyze the differences in the CTV dilution which is used as a measure of T_conv_ cell proliferation in response to co-culture with Tregs pre-treated with sgScramble supernatant or sgKdm6a supernatant.

### Isolation and enrichment of histones for liquid chromatography tandem mass spectrometry (LC–MS/MS)

Tregs were generated *in-vitro* as described above and then treated with or without 25 mM Na-La for 48 hours. Following treatment, histones were isolated using the Active Motif Histone Purification Mini Kit according to the manufacturer’s instructions. Briefly cells were harvested and washed by centrifugation at 200 x g for 5 minutes at 4°C. The resulting pellet was resuspended in pre-chilled extraction buffer and incubated overnight at 4°C with end-to-end rotation. The subsequent day, the samples were centrifuged at maximum speed for 5 minutes at 4°C and the supernatant containing the crude histones was then transferred to a fresh Eppendorf tube and neutralized with 5X Neutralization Buffer. The neutralized histone samples were then loaded onto a spin column and centrifuged at 2000 x g for 5 minutes at 4°C. The column was then washed thrice with Histone wash buffer and then the histones were eluted using the provided elution buffer, by centrifugation at 2000 x g for 5 minutes at 4°C and the histones were collected in the flow-through. For the precipitation of the eluted histones, perchloric acid was added to obtain a final concentration of 4% and the mixture was incubated overnight at 4°C. The next day, the samples were centrifuged at maximum speed at 4°C for an hour. The pelleted histones were then washed twice with cold 4% perchloric acid followed by two washes with ice-cold acetone (with 0.2% HCl) and pre-chilled acetone. The pellets were air-dried for 15 minutes and then resuspended in 100 mM tetraethylammonium bromide (pH 8) solutionby incubation for 15 minutes at room temperature with periodic vortexing.

The extracted histones were then quantitated using the Pierce BCA Assay Kit as per the manufacturer’s instructions and the absorbance was measured at 562 nm using the BioTek Synergy H1 micro-plate reader. 1000 µg of purified histones were then subjected to trypsin digestion overnight at 1:50 (trypsin to histone) ratio following the manufacturer’s guidelines. The digested samples were then reduced using 5 mM Dithiothreitol at 37°C for one hour and subsequently alkylated with 11 mM Iodoacetamide in the dark at room temperature for 50 minutes.

The tryptically digested histones were then immunoprecipitated using a Pan anti-Kla antibody using Pierce Classic IP Kit following the manufacturer’s protocol. 10 µg of Pan anti-Kla antibody was incubated overnight with 1000 µg of digested histone peptide solution at 4°C with end-to-end rotation. The next day, Pierce Protein A/G Agarose slurry solution was added into the Pierce Spin Column and centrifuged for 1 minute at 1000 × g. Then the antibody/ digested peptide mixture was added onto the Protein A/G Plus Agarose in the spin column and incubated over end-over-end rotation for 3.5 hours. The column was then washed thrice using 1X [conc] TBS Buffer and once with 1X Conditioning Buffer. The immune complex was eluted out of the column following incubation with low-pH elution buffer for 15 minutes and then centrifugation at 1000 x g for 1 minute at 4°C. The enriched peptides were then quantitated by BCA assay as described previously and sent to the MDACC Proteomics Core for LC–MS/MS analysis.

### LC–MS/MS Analysis

For the analysis of histone lactylation, vacuum-dried tryptic peptides were dissolved in solvent A (0.1% formic acid in water) and directly loaded onto a nano reverse-phase C18 column (75 μm ID × 25 cm L, 1.7 μm particle size). Peptides were separated using a linear gradient of solvent A (0.1% formic acid in water) and solvent B (0.1% formic acid in acetonitrile). The gradient was as follows: 0% to 10% solvent B over 2 minutes, 10% to 60% solvent B over 30 minutes, 60% to 100% solvent B over 1 minute, and held at 100% solvent B for 11 minutes. The eluate was electrosprayed into an Orbitrap Astral Mass Spectrometer operating in positive ion mode, with an electrospray voltage of 2 kV and an ion transfer tube temperature of 280°C. Data were acquired in DDA mode with a fixed cycle time of 2 seconds. Full MS scans were performed in the range of 350 to 1500 m/z at a resolution of 240,000, with an AGC target of 500% and a maximum injection time of 100 ms. Precursor ion selection width was set to 2 Th. Peptide fragmentation was carried out using HCD with a normalized collision energy of 27%. Fragment ion scans were recorded at a resolution of 80,000, with a maximum injection time of 30 ms and a normalized AGC target of 300%. Dynamic exclusion was enabled and set to 15 s.

For data analysis, the MS/MS spectra were processed using Proteome Discoverer v2.4 with a mouse protein sequence database from UniProt (downloaded November 2023, 17,184 entries). Peptide searching was performed with Sequest HT using the following parameters: allowing up to two missed cleavages, a precursor mass tolerance of 10 ppm, and a product ion tolerance of 0.6 Da. Carboxyamidomethylation of cysteine was set as a fixed modification, while oxidation of methionine and lactylation of lysine were set as variable modifications.

### ChIP-qPCR

*In-vitro* generated Tregs treated with and without 25mM Na-La for 48 hours and, *in-vitro* generated and CD4^+^CD25^+^ sorted Tregs treated with the supernatant of sgScramble or sgKdm6a cells for additional 24 hours were subjected to ChIP with the anti-H3K9la and anti-H3K18la antibody as previously described. For each condition, 100ng of immunoprecipitated DNA was used as input for real-time PCR (qPCR) to evaluate the gene expression utilizing the SYBR Green Master Mix according to the manufacturer’s instructions. For each condition, 100ng of immunoprecipitated DNA was used for real-time PCR (qPCR) to evaluate the gene expression, utilizing SYBR Green Master Mix according to the manufacturer’s instructions.

### RNA Isolation, cDNA preparation and qPCR

*In-vitro* generated Tregs treated with or without 25mM Na-La for 48 hours and, *in-vitro* generated and CD4^+^CD25^+^ sorted Tregs treated with the supernatant of sgScramble and sgKdm6a cells for 24 hours were lysed using the TriZol reagent and one-fifth volume of chloroform was added. After vigorous mixing and a 15 minute incubation at room temperature the samples were centrifuged at 12,000 X g for 15 minutes at 4°C. The aqueous layer on top was carefully collected, and incubated with equal volume of isopropanol for 10 minutes at room temperature for total RNA precipitation. Following this, the samples were centrifuged at 12,000 x g for 15 minutes at 4°C. The pellets were then washed with 75% ethanol by centrifugation. The resulting RNA pellets were air-dried, resuspended in nuclease-free water and quantified using a Nanodrop spectrophotometer. Subsequently, cDNA synthesis was performed by reverse transcribing 1µg of RNA using the Superscript III cDNA synthesis kit, according to the manufacturer’s instructions containing reverse transcriptase (RT) buffer, dNTP mix, RT Random primers, and Reverse transcriptase enzyme. 200ng of cDNA was then utilized for downstream qPCR analysis to measure gene expression using the SYBR Green Master Mix according to the manufacturer’s instructions.

## Data and code availability

The ChIP-seq and RNA-seq data supporting this study will be deposited in Sequence Read Archive (SRA). All requests for data should be made to the corresponding author, following verification of any intellectual property or confidentiality obligations.

This paper does not report original code.

Any additional information required to reanalyze the data reported in this paper will be available from the lead contact upon reasonable request.

## Ethics Statement

The research detailed in this study adheres to all relevant ethical regulations. Animal experiments were performed in accordance with the protocols approved by the Animal Resource Center at The University of Texas MD Anderson Cancer Center (UT MDACC).

## Acknowledgements

This research is supported by the Robert J. Kleberg, Jr. and Helen C. Kleberg Foundation Fund (FP00012831) (S.G), James P. Allison Institute Assistant Member Fund (S.G.), the MD Anderson Physician Scientist Award (S.G.), and NIH-R01836 Merit Award (R37 CA279192-01) (S.G.). We thank the MDACC Advanced Microscopy core (NIH grant S10 RR029552) for the comet assay and gamma-H2AX assay imaging services and the MDACC Metabolomics core (CCSG Grant NIH P30CA016672) for the Seahorse XFe96 Flux Analyzer service. We acknowledge the MDACC Proteomics core (UTMDACC NIH grants S10OD012304-01) for the LC-MS/MS study and the MDACC ATGC core (grant CA016672) for the sequencing studies. S.G. and P. Sharma are James P. Allison Institute members. P. Sharma is a member of the Parker Institute for Cancer Immunotherapy.

## Contributions

P.Singh and D.R. designed and performed the experiments, analyzed the data, and wrote the manuscript. B.C., S.M., and A.J.T. performed experiments and wrote the manuscript. A.M. and J.H. assisted with some *in vitro* experiments. Y.X. and P.L. performed the LC-MS/MS analysis. P. Sharma and P.P provided scientific input and edited the manuscript. S.G. developed the project, designed experiments, analyzed data, wrote the manuscript, and acquired funding.

## Supplemental information

**Figure S1.**
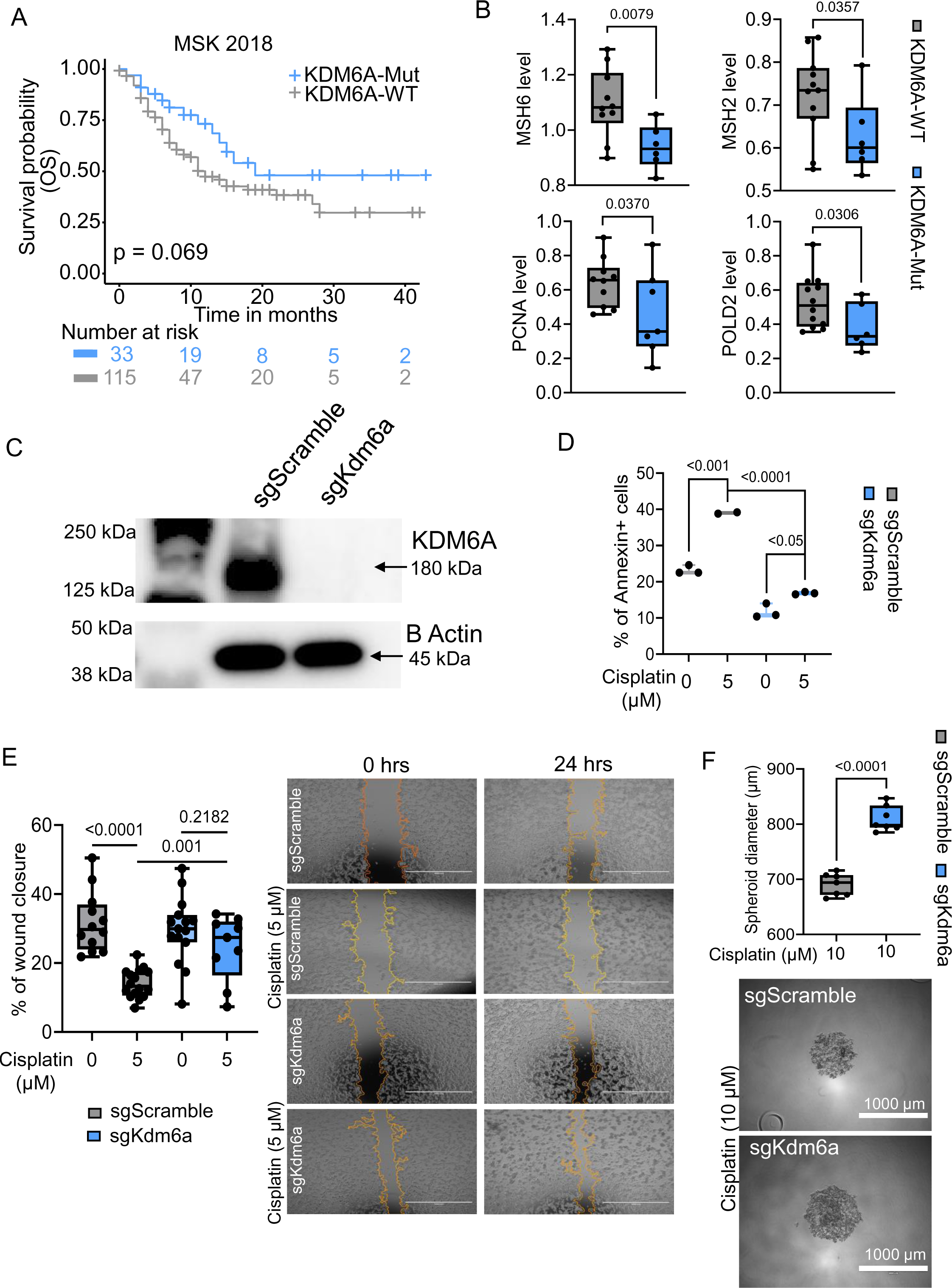
**(A)** Kaplan-Meier plots showing OS of KDM6A-WT and KDM6A-Mut patients receiving ICT, from MSK_2018 cohort (n=148 patients, KDM6A-Mut=33, KDM6A-WT=115). Two-tailed Log-rank test was performed. **(B)** Box-and-whisker plots depicting the expression level of indicated MMR pathway associated genes between KDM6A-WT and KDM6A-Mut patients in HCRN cohort. One-tailed Student’s t-test was performed. **(C)** Representative western blot image indicating the expression of KDM6A (180 kDa) and Housekeeping β-ACTIN (45 kDa) in sgScramble and sgKdm6a cells lines. Data is representative of two independent experiments. **(D)** Box-and-whisker plot demonstrating percentage of Annexin+ sgScramble and sgKdm6a cells treated with or without 5 µM cisplatin for 48 hours. (n=3 biologically independent samples per group) Two-tailed Student’s t-test was performed. **(E)** Box-and-whisker plot (Left) and representative images (Right) depicting the percentage of wound closure of sgScramble and sgKdm6a cells treated with or without 5 µM cisplatin for 24 hours. (n=9 biologically independent samples per group) Two-tailed Student’s t-test was performed. For the representative images, data is indicative of two independent experiments. **(F)** Box-and-whisker plot (Left) and representative images (Right) showing the diameter of sgScramble and sgKdm6a spheroids (n=7 biologically independent samples per group). Two-tailed Student’s t-test was performed. For the representative images, data is indicative of two independent experiments. For all the box-and-whisker plots in Figure S1, the centre line marks the median, the edges of the box represent the interquartile (25th–75th) percentile and the whiskers represent minimum-maximum values.

**Figure S2.**
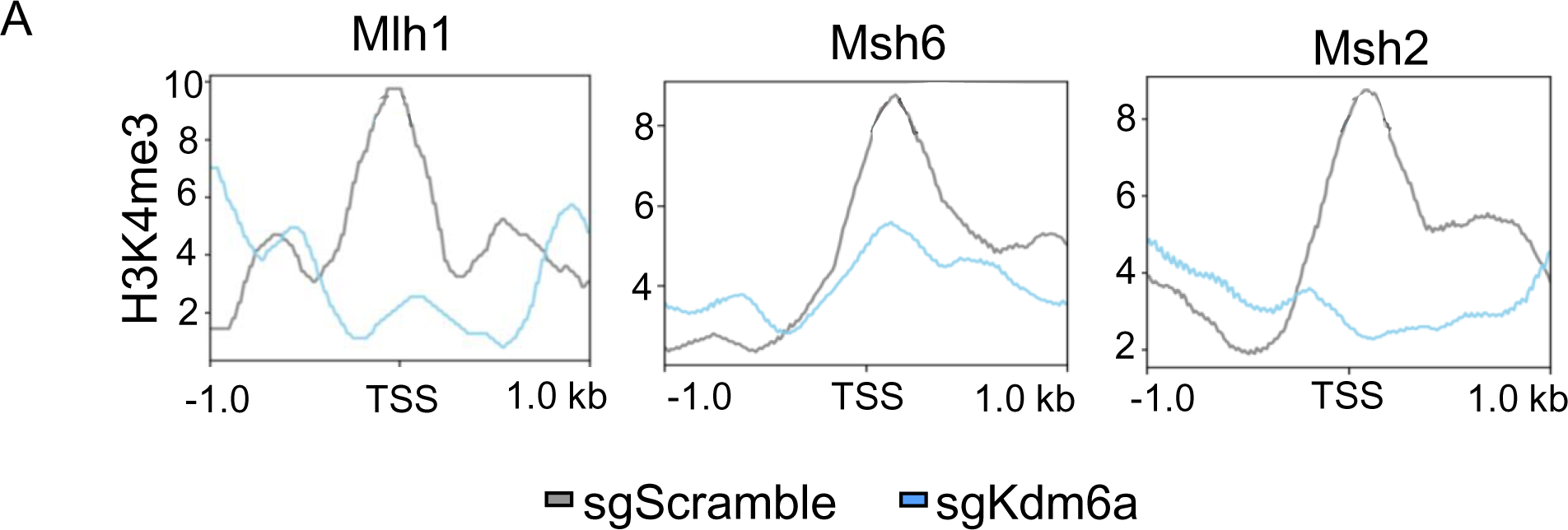
**(A)** Profile plots depicting the probability scores of H3K4me3 binding at promoter regions (TSS ±1kb) of *Mlh1*, *Msh6* and *Msh2* gene loci in sgScramble and sgKdm6a cell lines.

**Figure S3.**
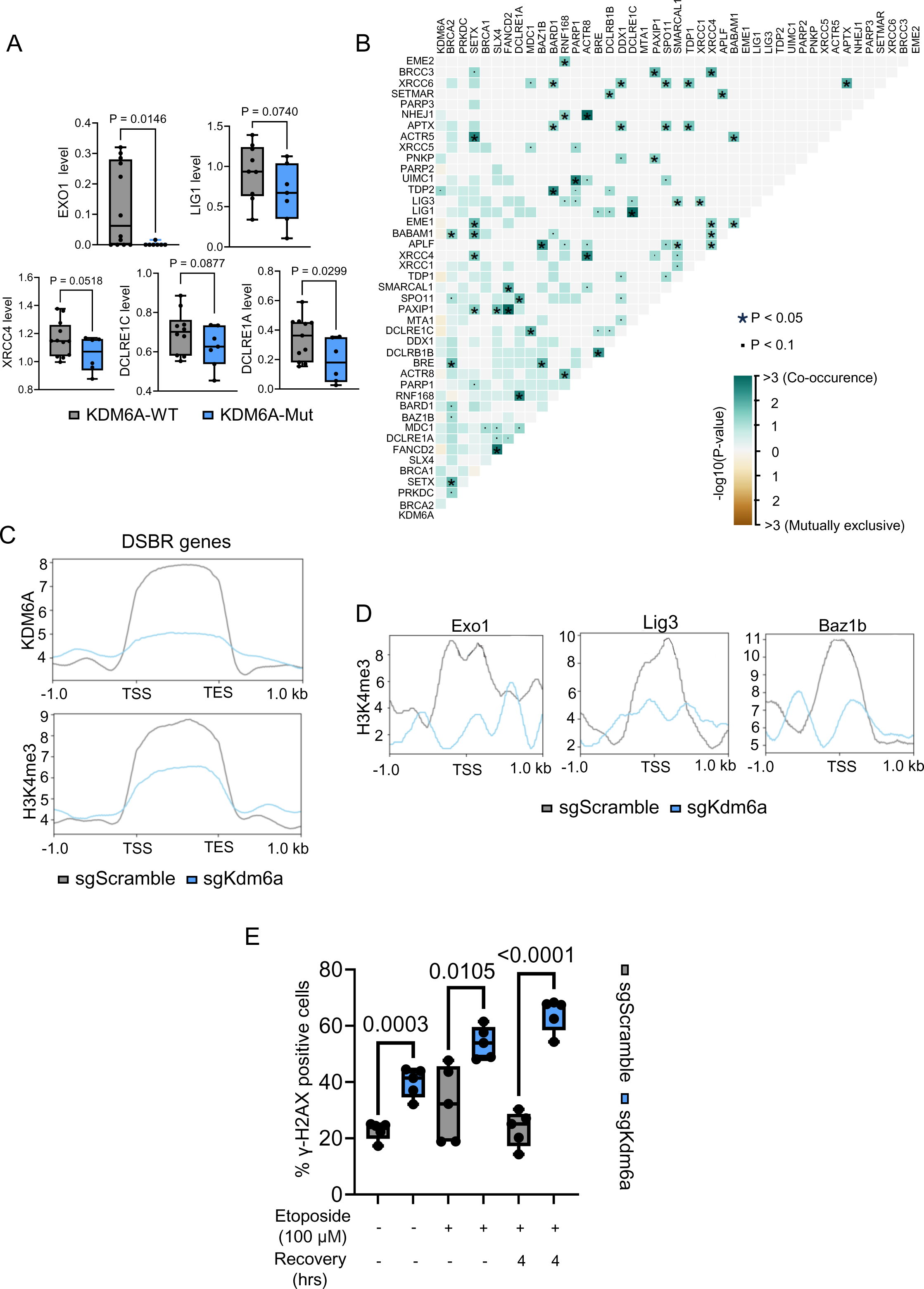
**(A)** Box-and-whisker plots depicting expression level of indicated DSBR genes between KDM6A-WT and KDM6A-Mut patients in HCRN cohort. One-tailed Student’s t-test was performed. **(B)** Correlation plot displaying co-occurrence or mutual exclusivity of *KDM6A* mutation with DSBR gene mutations in bladder cancer patients in the TCGA BLCA (n=412) dataset harboring *KDM6A* mutation. Two-tailed Pair wise Fisher’s exact test was performed. **(C)** Profile plots representing the probability score of KDM6A and H3K4me3 binding at ±1kb regions from TSS and TES for genes involved in DSBR pathway in sgScramble and sgKdm6a cell lines. **(D)** Profile plots showing the probability scores of H3K4me3 binding at promoter regions (TSS ±1kb) of *Exo1, Lig3 and Baz1b* gene loci in sgScramble and sgKdm6a cell lines. **(E)** Box-and-whisker plot indicating the percentage of gamma-H2AX+ sgScramble and sgKdm6a cells treated with and without 100µM Etoposide followed by 4 hours of recovery or no recovery (n=5 biologically independent samples per group). Two-tailed Student’s t-test was performed. For the box-and-whisker plot in Figure S3, the centre line marks the median, the edges of the box represent the interquartile (25th–75th) percentile and the whiskers represent minimum-maximum values.

**Figure S4.**
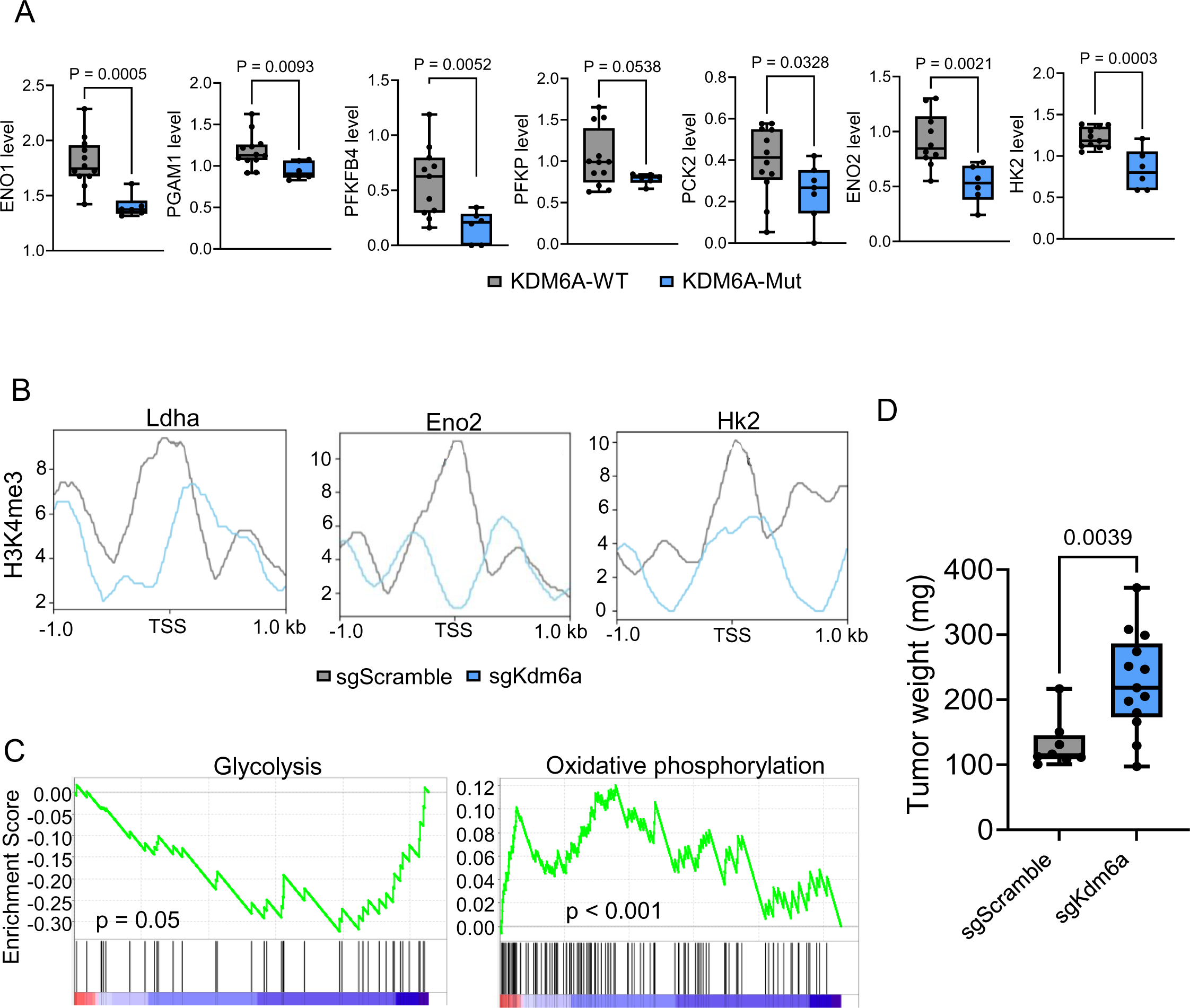
**(A)** Box-and-whisker plots showing expression level of glycolytic genes between KDM6A-WT and KDM6A-Mut in in HCRN cohort. One-tailed Student’s t-test was performed. **(B)** Profile plots indicating the probability scores of H3K4me3 binding at promoter regions (TSS ±1kb) of *Ldha, Hk2 and Eno2* gene loci in sgScramble and sgKdm6a cell lines. **(C)** Plots representing GSEA pathways generated from genes with KDM6A enrichment in sgKdm6a cells compared to sgScramble cells. Indicated P-values represent one-tailed nominal p-value of gene enrichments calculated by permutations. **(D)** Box-and-whisker plots demonstrating the weights of sgScramble and sgKdm6a tumor bearing female mice (n=8 sgScramble and 13 sgKdm6a female tumor bearing mice). For all the box-and-whisker plots in Figure S4, the centre line marks the median, the edges of the box represent the interquartile (25th–75th) percentile and the whiskers represent minimum-maximum values.

**Figure S5.**
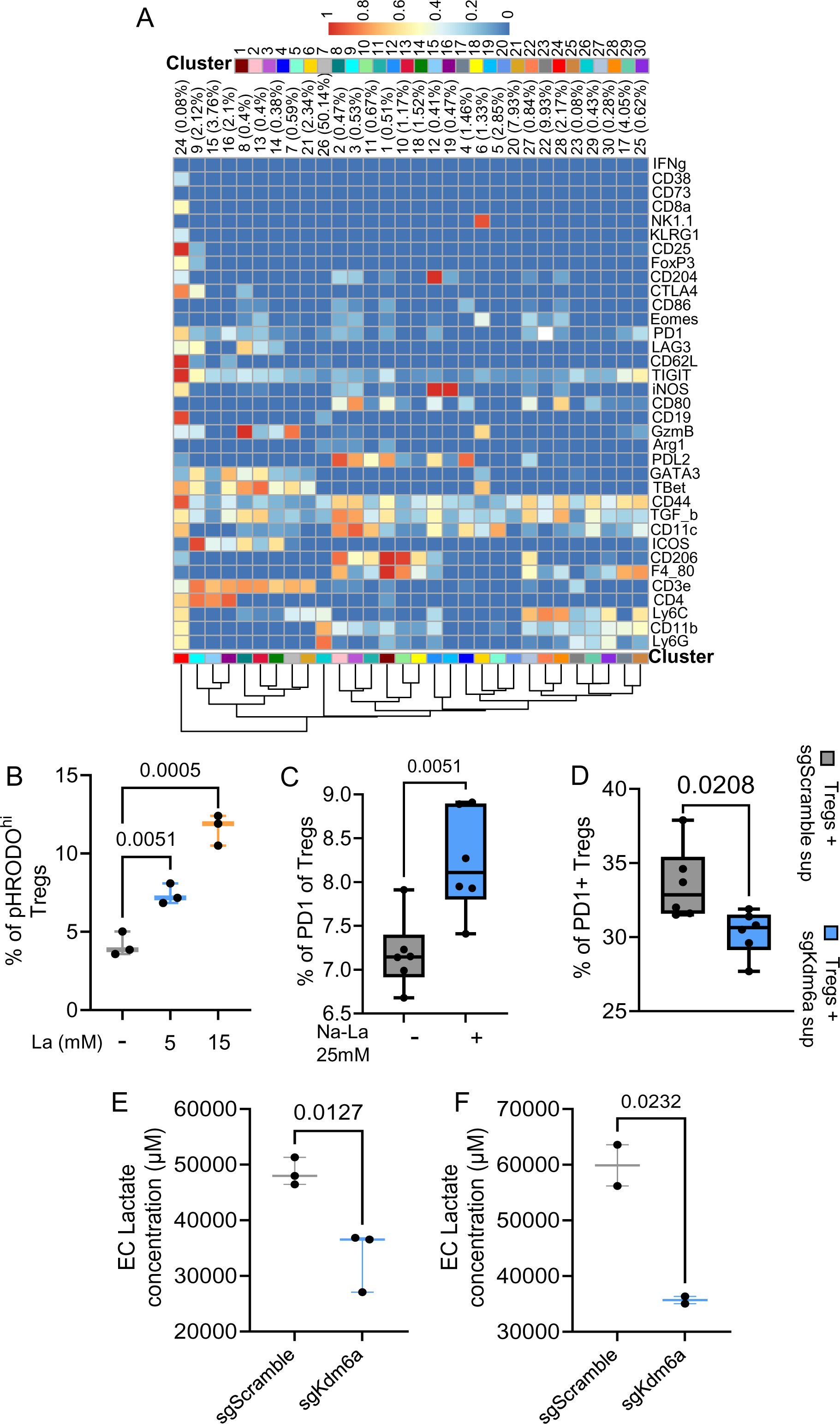
**(A)** Heatmap indicating the expression of genes of interest in the different CD45^+^ immune cell clusters (shown in Figure 5A) as determined by mass cytometry. **(B)** Box-and-whisker plot representing the percentage of pHRODO^hi^ cells in *in-vitro* generated Tregs treated with or without 5mM and 15mM Lactic acid (La) for 30 minutes (n=3 biologically independent samples per group). Two-tailed Student’s t test was performed. **(C)** Box-and-whisker plot demonstrating the percentage of PD-1^+^ cells in *in-vitro* generated Tregs treated with and without 25 mM Na-La for 48 hours (n=6 biologically independent samples per group). Two-tailed Student’s t test was performed. **(D)** Box-and-whisker plot showing the percentage of PD-1^+^ cells in *in-vitro* generated Tregs treated with the culture supernatants from sgScramble or sgKdm6a cell lines for 24 hours (n=6 biologically independent samples per group). Two-tailed Student’s t-test was performed. **(E)** Box-and-whisker plot depicting the EC L-Lactate concentration in the supernatant of sgScramble and SgKdm6a cells cultured for 24 hours (n=3 biologically independent samples per group). These supernatants were subsequently utilized to treat the Tregs in Figure 5H and S5D. Two-tailed Student’s t-test was performed. **(F)** Box-and-whisker plot displaying the EC L-Lactate concentration in the supernatant of sgScramble and SgKdm6a cell lines cultured for 36 hours (n=2 biologically independent samples per group). These supernatants were subsequently utilized to treat the Tregs in Figure 5I. Two-tailed Student’s t-test was performed. For all the box-and-whisker plots in Figure S5, the centre line marks the median, the edges of the box represent the interquartile (25th–75th) percentile and the whiskers represent minimum-maximum values.

**Figure S6.**
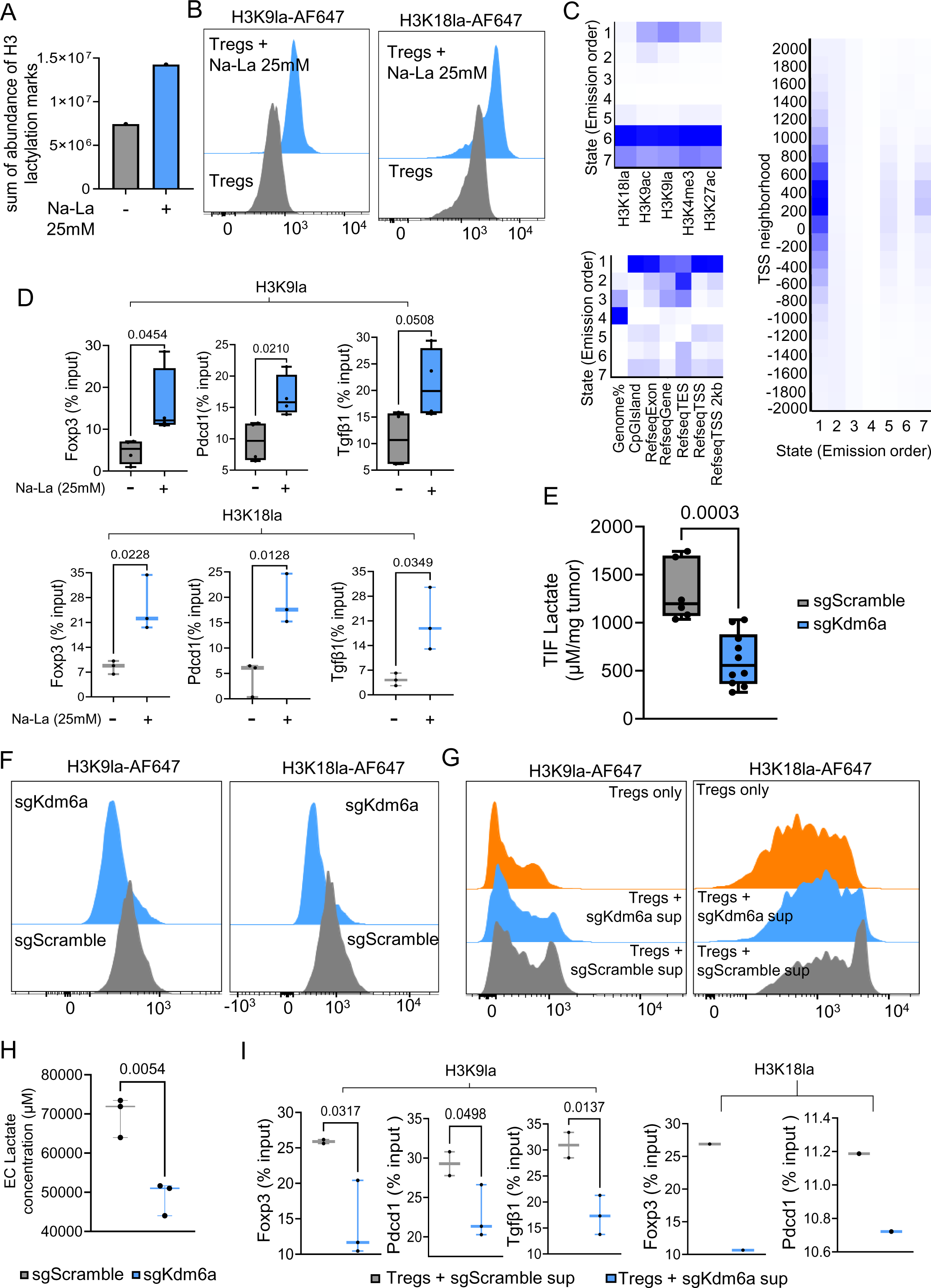
**(A)** Box-and-whisker plot illustrating the sum of abundance of H3 lactylation marks in *in-vitro* generated Tregs treated with and without 25 mM Na-La for 48 hours as determined by liquid chromatography with tandem mass spectrometry (LC–MS/MS). **(B)** Representative histogram depicting the MFI of H3K9la (Left Panel) and H3K18la (Right Panel) in *in-vitro* generated Tregs treated with and without 25 mM Na-La for 48 hours. **(C)** Heatmap indicating emission parameters, generated from ChromHMM analysis. Each row corresponds to a specific chromatin state and each column corresponds to a specific epigenetic mark in *in-vitro* generated Tregs (Top left Panel). Heatmap displaying the fold enrichment of different chromatin states in distinct genomic regions based on the indicated epigenetic marks in *in-vitro* generated Tregs (Bottom left Panel). Heatmap demonstrating the fold enrichment of different chromatin states around transcription start sites (TSS) based on indicated epigenetic marks, in *in-vitro* generated Tregs (Right panel). The darker blue color corresponds to a greater probability of observing the mark in the state. **(D)** Box-and-whisker plots demonstrating the ChIP qPCR analysis of H3K9la (Top Panel) and H3K18la (Bottom Panel) enrichment at the indicated gene promoters in *in-vitro* generated Tregs treated with and without 25 mM Na-La for 48 hours. Data is represented as percentage of input (n=4 (Top) and 3 (Bottom) biologically independent samples per group). Two tailed Student’s t-test was performed. **(E)** Box-and-whisker plot showing the L-lactate concentrations in the TIF normalized to corresponding tumor weights, obtained from sgScramble and sgKdm6a tumors (n=6 sgScramble and 10 sgKdm6a tumor bearing mice). Two-tailed Student’s t test was performed. **(F)** Representative histogram displaying the MFI of H3K9la (Left Panel) and H3K18la (Right Panel) in intratumoral Tregs derived from sgScramble and sgKdm6a tumor harboring mice. **(G)** Representative histogram representing the MFI of H3K9la (Left Panel) and H3K18la (Right Panel) in *in-vitro* generated Tregs treated with and without culture supernatants from sgScramble and sgKdm6a cell lines for 24 hours. **(H)** Box-and-whisker plot representing the Extracellular (EC) L-Lactate concentration in the supernatants of sgScramble and sgKdm6a cell lines, measured following 24 hours of culture (n=3 biologically independent samples per group). These supernatants were subsequently utilized to treat the Tregs in Figure 6F, 6G, S6I.Two-tailed Student’s t-test was performed. **(I)** Box-and-whisker plots showing the ChIP qPCR analysis of H3K9la and H3K18la enrichment at the indicated gene promoters in *in-vitro* generated Tregs treated with culture supernatants from sgScramble and sgKdm6a cell lines for 24 hours (n=3). One tailed Student’s t-test was performed. For all the box-and-whisker plots in Figure S6, the centre line marks the median, the edges of the box represent the interquartile (25th–75th) percentile and the whiskers represent minimum-maximum values.

## References

1. Siegel RL, Giaquinto AN, Jemal A. Cancer statistics, 2024. CA Cancer J Clin. 2024;74(1):12–49.

2. Alifrangis C, McGovern U, Freeman A, Powles T, Linch M. Molecular and histopathology directed therapy for advanced bladder cancer. Nat Rev Urol. 2019;16(8):465–83.

3. Felsenstein KM, Theodorescu D. Precision medicine for urothelial bladder cancer: update on tumour genomics and immunotherapy. Nat Rev Urol. 2018;15(2):92–111.

4. Robertson AG, Kim J, Al-Ahmadie H, Bellmunt J, Guo G, Cherniack AD, et al. Comprehensive Molecular Characterization of Muscle-Invasive Bladder Cancer. Cell. 2018;174(4):1033.

5. Wolff EM, Liang G, Jones PA. Mechanisms of Disease: genetic and epigenetic alterations that drive bladder cancer. Nat Clin Pract Urol. 2005;2(10):502–10.

6. Baylin SB, Jones PA. A decade of exploring the cancer epigenome - biological and translational implications. Nat Rev Cancer. 2011;11(10):726–34.

7. Tran L, Xiao JF, Agarwal N, Duex JE, Theodorescu D. Advances in bladder cancer biology and therapy. Nat Rev Cancer. 2021;21(2):104–21.

8. Qiu H, Makarov V, Bolzenius JK, Halstead A, Parker Y, Wang A, et al. KDM6A Loss Triggers an Epigenetic Switch That Disrupts Urothelial Differentiation and Drives Cell Proliferation in Bladder Cancer. Cancer Res. 2023;83(6):814–29.

9. Rachakonda PS, Hosen I, de Verdier PJ, Fallah M, Heidenreich B, Ryk C, et al. TERT promoter mutations in bladder cancer affect patient survival and disease recurrence through modification by a common polymorphism. Proc Natl Acad Sci U S A. 2013;110(43):17426–31.

10. Glaser AP, Fantini D, Shilatifard A, Schaeffer EM, Meeks JJ. The evolving genomic landscape of urothelial carcinoma. Nat Rev Urol. 2017;14(4):215–29.

11. Hurst CD, Alder O, Platt FM, Droop A, Stead LF, Burns JE, et al. Genomic Subtypes of Non-invasive Bladder Cancer with Distinct Metabolic Profile and Female Gender Bias in KDM6A Mutation Frequency. Cancer Cell. 2017;32(5):701–15 e7.

12. Agger K, Cloos PA, Christensen J, Pasini D, Rose S, Rappsilber J, et al. UTX and JMJD3 are histone H3K27 demethylases involved in HOX gene regulation and development. Nature. 2007;449(7163):731-4.

13. Lan F, Bayliss PE, Rinn JL, Whetstine JR, Wang JK, Chen S, et al. A histone H3 lysine 27 demethylase regulates animal posterior development. Nature. 2007;449(7163):689–94.

14. Mariathasan S, Turley SJ, Nickles D, Castiglioni A, Yuen K, Wang Y, et al. TGFbeta attenuates tumour response to PD-L1 blockade by contributing to exclusion of T cells. Nature. 2018;554(7693):544–8.

15. Samstein RM, Lee CH, Shoushtari AN, Hellmann MD, Shen R, Janjigian YY, et al. Tumor mutational load predicts survival after immunotherapy across multiple cancer types. Nat Genet. 2019;51(2):202–6.

16. Pietzak EJ, Zabor EC, Bagrodia A, Armenia J, Hu W, Zehir A, et al. Genomic Differences Between “Primary” and “Secondary” Muscle-invasive Bladder Cancer as a Basis for Disparate Outcomes to Cisplatin-based Neoadjuvant Chemotherapy. Eur Urol. 2019;75(2):231–9.

17. Clinton TN, Chen Z, Wise H, Lenis AT, Chavan S, Donoghue MTA, et al. Genomic heterogeneity as a barrier to precision oncology in urothelial cancer. Cell Rep. 2022;41(12):111859.

18. von Loga K, Woolston A, Punta M, Barber LJ, Griffiths B, Semiannikova M, et al. Extreme intratumour heterogeneity and driver evolution in mismatch repair deficient gastro-oesophageal cancer. Nat Commun. 2020;11(1):139.

19. Damrauer JS, Beckabir W, Klomp J, Zhou M, Plimack ER, Galsky MD, et al. Collaborative study from the Bladder Cancer Advocacy Network for the genomic analysis of metastatic urothelial cancer. Nat Commun. 2022;13(1):6658.

20. Lipkin SM, Wang V, Jacoby R, Banerjee-Basu S, Baxevanis AD, Lynch HT, et al. MLH3: a DNA mismatch repair gene associated with mammalian microsatellite instability. Nat Genet. 2000;24(1):27–35.

21. Chung J, Maruvka YE, Sudhaman S, Kelly J, Haradhvala NJ, Bianchi V, et al. DNA Polymerase and Mismatch Repair Exert Distinct Microsatellite Instability Signatures in Normal and Malignant Human Cells. Cancer Discov. 2021;11(5):1176–91.

22. Maio M, Ascierto PA, Manzyuk L, Motola-Kuba D, Penel N, Cassier PA, et al. Pembrolizumab in microsatellite instability high or mismatch repair deficient cancers: updated analysis from the phase II KEYNOTE-158 study. Ann Oncol. 2022;33(9):929–38.

23. Grothey A. Pembrolizumab in MSI-H-dMMR Advanced Colorectal Cancer - A New Standard of Care. N Engl J Med. 2020;383(23):2283–5.

24. Aebi S, Kurdi-Haidar B, Gordon R, Cenni B, Zheng H, Fink D, et al. Loss of DNA mismatch repair in acquired resistance to cisplatin. Cancer Res. 1996;56(13):3087–90.

25. Silva MM, Rocha CRR, Kinker GS, Pelegrini AL, Menck CFM. The balance between NRF2/GSH antioxidant mediated pathway and DNA repair modulates cisplatin resistance in lung cancer cells. Sci Rep. 2019;9(1):17639.

26. Sawant A, Kothandapani A, Zhitkovich A, Sobol RW, Patrick SM. Role of mismatch repair proteins in the processing of cisplatin interstrand cross-links. DNA Repair (Amst). 2015;35:126–36.

27. Tuninetti V, Pace L, Ghisoni E, Quara V, Arezzo F, Palicelli A, et al. Retrospective Analysis of the Correlation of MSI-h/dMMR Status and Response to Therapy for Endometrial Cancer: RAME Study, a Multicenter Experience. Cancers (Basel). 2023;15(14).

28. Olive PL, Banath JP. The comet assay: a method to measure DNA damage in individual cells. Nat Protoc. 2006;1(1):23–9.

29. Chatzidoukaki O, Goulielmaki E, Schumacher B, Garinis GA. DNA Damage Response and Metabolic Reprogramming in Health and Disease. Trends Genet. 2020;36(10):777–91.

30. Milanese C, Bombardieri CR, Sepe S, Barnhoorn S, Payan-Gomez C, Caruso D, et al. DNA damage and transcription stress cause ATP-mediated redesign of metabolism and potentiation of anti-oxidant buffering. Nat Commun. 2019;10(1):4887.

31. Sobanski T, Rose M, Suraweera A, O’Byrne K, Richard DJ, Bolderson E. Cell Metabolism and DNA Repair Pathways: Implications for Cancer Therapy. Front Cell Dev Biol. 2021;9:633305.

32. De Martino M, Rathmell JC, Galluzzi L, Vanpouille-Box C. Cancer cell metabolism and antitumour immunity. Nat Rev Immunol. 2024.

33. Watson MJ, Vignali PDA, Mullett SJ, Overacre-Delgoffe AE, Peralta RM, Grebinoski S, et al. Metabolic support of tumour-infiltrating regulatory T cells by lactic acid. Nature. 2021;591(7851):645–51.

34. Batlle E, Massague J. Transforming Growth Factor-beta Signaling in Immunity and Cancer. Immunity. 2019;50(4):924–40.

35. Zhang D, Tang Z, Huang H, Zhou G, Cui C, Weng Y, et al. Metabolic regulation of gene expression by histone lactylation. Nature. 2019;574(7779):575–80.

36. Raychaudhuri D, Singh P, Hennessey M, Chakraborty B, Tannir AJ, Trujillo-Ocampo A, et al. Histone Lactylation Drives CD8 T Cell Metabolism and Function. bioRxiv. 2024.

37. Dyrskjot L, Hansel DE, Efstathiou JA, Knowles MA, Galsky MD, Teoh J, et al. Bladder cancer. Nat Rev Dis Primers. 2023;9(1):58.

38. Robertson AG, Kim J, Al-Ahmadie H, Bellmunt J, Guo G, Cherniack AD, et al. Comprehensive Molecular Characterization of Muscle-Invasive Bladder Cancer. Cell. 2017;171(3):540–56 e25.

39. Chen X, Lin X, Pang G, Deng J, Xie Q, Zhang Z. Significance of KDM6A mutation in bladder cancer immune escape. BMC Cancer. 2021;21(1):635.

40. Akkers RC, van Heeringen SJ, Jacobi UG, Janssen-Megens EM, Francoijs KJ, Stunnenberg HG, et al. A hierarchy of H3K4me3 and H3K27me3 acquisition in spatial gene regulation in Xenopus embryos. Dev Cell. 2009;17(3):425–34.

41. Lim PS, Li J, Holloway AF, Rao S. Epigenetic regulation of inducible gene expression in the immune system. Immunology. 2013;139(3):285–93.

42. Leng X, Wang J, An N, Wang X, Sun Y, Chen Z. Histone 3 lysine-27 demethylase KDM6A coordinates with KMT2B to play an oncogenic role in NSCLC by regulating H3K4me3. Oncogene. 2020;39(41):6468–79.

43. Chen Z, Qi Y, Shen J, Chen Z. Histone demethylase KDM6A coordinating with KMT2B regulates self-renewal and chemoresistance of non-small cell lung cancer stem cells. Transl Oncol. 2023;37:101778.

44. Boila LD, Ghosh S, Bandyopadhyay SK, Jin L, Murison A, Zeng AGX, et al. KDM6 demethylases integrate DNA repair gene regulation and loss of KDM6A sensitizes human acute myeloid leukemia to PARP and BCL2 inhibition. Leukemia. 2023;37(4):751–64.

45. Lee JM, Ledermann JA, Kohn EC. PARP Inhibitors for BRCA1/2 mutation-associated and BRCA-like malignancies. Ann Oncol. 2014;25(1):32–40.

46. Farmer H, McCabe N, Lord CJ, Tutt AN, Johnson DA, Richardson TB, et al. Targeting the DNA repair defect in BRCA mutant cells as a therapeutic strategy. Nature. 2005;434(7035):917–21.

47. Lobo N, Mount C, Omar K, Nair R, Thurairaja R, Khan MS. Landmarks in the treatment of muscle-invasive bladder cancer. Nat Rev Urol. 2017;14(9):565–74.

48. Galluzzi L, Senovilla L, Vitale I, Michels J, Martins I, Kepp O, et al. Molecular mechanisms of cisplatin resistance. Oncogene. 2012;31(15):1869–83.

49. Tan Y, Li J, Zhao G, Huang KC, Cardenas H, Wang Y, et al. Metabolic reprogramming from glycolysis to fatty acid uptake and beta-oxidation in platinum-resistant cancer cells. Nat Commun. 2022;13(1):4554.

50. Xu T, Junge JA, Delfarah A, Lu YT, Arnesano C, Iqbal M, et al. Bladder cancer cells shift rapidly and spontaneously to cisplatin-resistant oxidative phosphorylation that is trackable in real time. Sci Rep. 2022;12(1):5518.

51. Gu J, Zhou J, Chen Q, Xu X, Gao J, Li X, et al. Tumor metabolite lactate promotes tumorigenesis by modulating MOESIN lactylation and enhancing TGF-beta signaling in regulatory T cells. Cell Rep. 2022;39(12):110986.

52. Kumagai S, Koyama S, Itahashi K, Tanegashima T, Lin YT, Togashi Y, et al. Lactic acid promotes PD-1 expression in regulatory T cells in highly glycolytic tumor microenvironments. Cancer Cell. 2022;40(2):201–18 e9.

53. Langmead B, Salzberg SL. Fast gapped-read alignment with Bowtie 2. Nat Methods. 2012;9(4):357–9.

54. Faust GG, Hall IM. SAMBLASTER: fast duplicate marking and structural variant read extraction. Bioinformatics. 2014;30(17):2503–5.

55. Li H, Handsaker B, Wysoker A, Fennell T, Ruan J, Homer N, et al. The Sequence Alignment/Map format and SAMtools. Bioinformatics. 2009;25(16):2078–9.

56. Tarasov A, Vilella AJ, Cuppen E, Nijman IJ, Prins P. Sambamba: fast processing of NGS alignment formats. Bioinformatics. 2015;31(12):2032–4.

57. Zhang Y, Liu T, Meyer CA, Eeckhoute J, Johnson DS, Bernstein BE, et al. Model-based analysis of ChIP-Seq (MACS). Genome Biol. 2008;9(9):R137.

58. Wang Q, Li M, Wu T, Zhan L, Li L, Chen M, et al. Exploring Epigenomic Datasets by ChIPseeker. Curr Protoc. 2022;2(10):e585.

59. Yu G, Wang LG, Han Y, He QY. clusterProfiler: an R package for comparing biological themes among gene clusters. OMICS. 2012;16(5):284–7.

60. Shao Z, Zhang Y, Yuan GC, Orkin SH, Waxman DJ. MAnorm: a robust model for quantitative comparison of ChIP-Seq data sets. Genome Biol. 2012;13(3):R16.

61. Subramanian A, Tamayo P, Mootha VK, Mukherjee S, Ebert BL, Gillette MA, et al. Gene set enrichment analysis: a knowledge-based approach for interpreting genome-wide expression profiles. Proc Natl Acad Sci U S A. 2005;102(43):15545–50.

62. Ramirez F, Ryan DP, Gruning B, Bhardwaj V, Kilpert F, Richter AS, et al. deepTools2: a next generation web server for deep-sequencing data analysis. Nucleic Acids Res. 2016;44(W1):W160–5.

63. Ernst J, Kellis M. Chromatin-state discovery and genome annotation with ChromHMM. Nat Protoc. 2017;12(12):2478–92.

